# *F*_ST_ and kinship for arbitrary population structures II: Method-of-moments estimators

**DOI:** 10.1101/083923

**Authors:** Alejandro Ochoa, John D. Storey

## Abstract

*F*_ST_ and kinship are key parameters often estimated in modern population genetics studies in order to quantitatively characterize structure and relatedness. Kinship matrices have also become a fundamental quantity used in genome-wide association studies and heritability estimation. The most frequently used estimators of *F*_ST_ and kinship are method-of-moments estimators whose accuracies depend strongly on the existence of simple underlying forms of structure, such as the independent subpopulations model of non-overlapping, independently evolving subpopulations. However, modern data sets have revealed that these simple models of structure likely do not hold in many populations, including humans. In this work, we provide new results on the behavior of these estimators in the presence of arbitrarily complex population structures, which results in an improved estimation framework specifically designed for arbitrary population structures. After establishing a framework for assessing bias and consistency of genome-wide estimators, we calculate the accuracy of existing *F*_ST_ and kinship estimators under arbitrary population structures, characterizing biases and estimation challenges unobserved under their originally assumed models of structure. We then present our new approach, which consistently estimates kinship and *F*_ST_ when the minimum kinship value in the dataset is estimated consistently. We illustrate our results using simulated genotypes from an admixture model, constructing a one-dimensional geographic scenario that departs nontrivially from the independent subpopulations model. Our simulations reveal the potential for severe biases in estimates of existing approaches that are overcome by our new framework. This work may significantly improve future analyses that rely on accurate kinship and *F*_ST_ estimates.

## 1 Introduction

In population genetics studies, one is often interested in characterizing structure, genetic differentiation, and relatedness among individuals. Two quantities often considered in this context are *F*_ST_ and kinship. *F*_ST_ is a parameter that measures structure in a subdivided population, satisfying *F*_ST_ = 0 for an unstructured population and *F*_ST_ = 1 if every locus has become fixed for some allele in each subpopulation. More generally, *F*_ST_ is the probability that alleles drawn randomly from a subpopulation are “identical by descent” (IBD) relative to an ancestral population [3, 4]. The kinship coefficient is a measure of relatedness between individuals defined in terms of IBD probabilities, and it is closely related to *F*_ST_ [3].

This work focuses on the estimation of *F*_ST_ and kinship from biallelic single-nucleotide polymorphism (SNP) marker data. Existing estimators can be classified into parametric estimators (methods that require a likelihood function) and non-parametric estimators (such as the method-of-moments estimators we focus on, which only require low-order moment equations). There are many likelihood approaches that estimate *F*_ST_ and kinship, but these are limited by assuming independent subpopulations or Normal approximations for *F*_ST_ [5–13] or outbred individuals for kinship [14, 15]. Additionally, more complete likelihood models such as that of [16] are underdetermined for biallelic loci [17]. Non-parametric approaches such as those based on the method of moments are considerably more flexible and computationally tractable [18], so they are the natural choice to study arbitrary population structures.

The most frequently-used *F*_ST_ estimators are derived and justified under the “independent sub-populations model,” in which non-overlapping subpopulations evolved independently by splitting all at the same time from a common ancestral population. The Weir-Cockerham (WC) *F*_ST_ estimator assumes subpopulations of differing sample sizes and equal per-subpopulation *F*_ST_ relative to the common ancestral population [19]. The “Hudson” *F*_ST_ estimator [20] assumes two subpopulations with different *F*_ST_ values. These *F*_ST_ estimators are ratio estimators derived using the method of moments to have unbiased numerators and denominators, which gives approximately unbiased ratio estimates when their assumptions are met [6, 19, 20]. We also evaluate BayeScan [12], which estimates population-specific *F*_ST_ values using a Bayesian model and the Dirichlet-Multinomial likelihood function—thus representing non-method-of-moments approaches—but which like the WC and Hudson *F*_ST_ estimators also assumes that subpopulations are non-overlapping and evolve independently. These *F*_ST_ estimators are important contributions, used widely in the field.

Kinship coefficients are now commonly calculated in population genetics studies to capture structure and relatedness. Kinship is utilized in principal components analyses and linear-mixed effects models to correct for structure in Genome-Wide Association Studies (GWAS) [18, 21–27] and to estimate genome-wide heritability [28, 29]. Often absent in previous works is a clear identification and role of the ancestral population *T* that sets the scale of the kinship estimates used. Omission of *T* makes sense when kinship is estimated on an unstructured population (where only a few individual pairs are closely related; there *T* is the current population). Our more complete notation brings *T* to the fore and highlights its key role in kinship estimation and its applications. The most commonly-used kinship estimator [18, 24, 27–33] is also a method-of-moments estimator whose operating characteristics are largely unknown in the presence of structure. We show in Section 4 that this popular estimator is accurate only when the average kinship is zero, which implies that the population must be unstructured.

Recent genome-wide studies have revealed that humans and other natural populations are structured in a complex manner that break the assumptions of the above estimators. Such complex population structures has been observed in several large human studies, such as the Human Genome Diversity Project [34, 35], the 1000 Genomes Project [36], Human Origins [37–39], and other contemporary [40–44] and archaic populations [45, 46]. We have also demonstrated, based on the work in Part I and Part II here, that the global human population has a complex kinship matrix and no independent subpopulations [47]. Therefore, there is a need for innovative approaches designed for complex population structures. To this end, we reveal the operating characteristics of these frequently-used *F*_ST_ and kinship estimators in the presence of arbitrary forms of structure, which leads to a new estimation strategy for *F*_ST_ and kinship.

We generalized the definition of *F*_ST_ for arbitrary population structures in Section 3 of Part I. Additionally, we derived connections between *F*_ST_ and three models: arbitrary kinship coefficients [3, 16] in Section 3 of Part I (panel “Kinship Model” in Fig. 1), individual-specific allele frequencies [48, 49] in Section 5 of Part I (panels “Coancestry Model” and “Coancestry in Terms of Kinship” in Fig. 1), and admixture models [50–52] in Section 6 of Part I.

**Figure 1.**
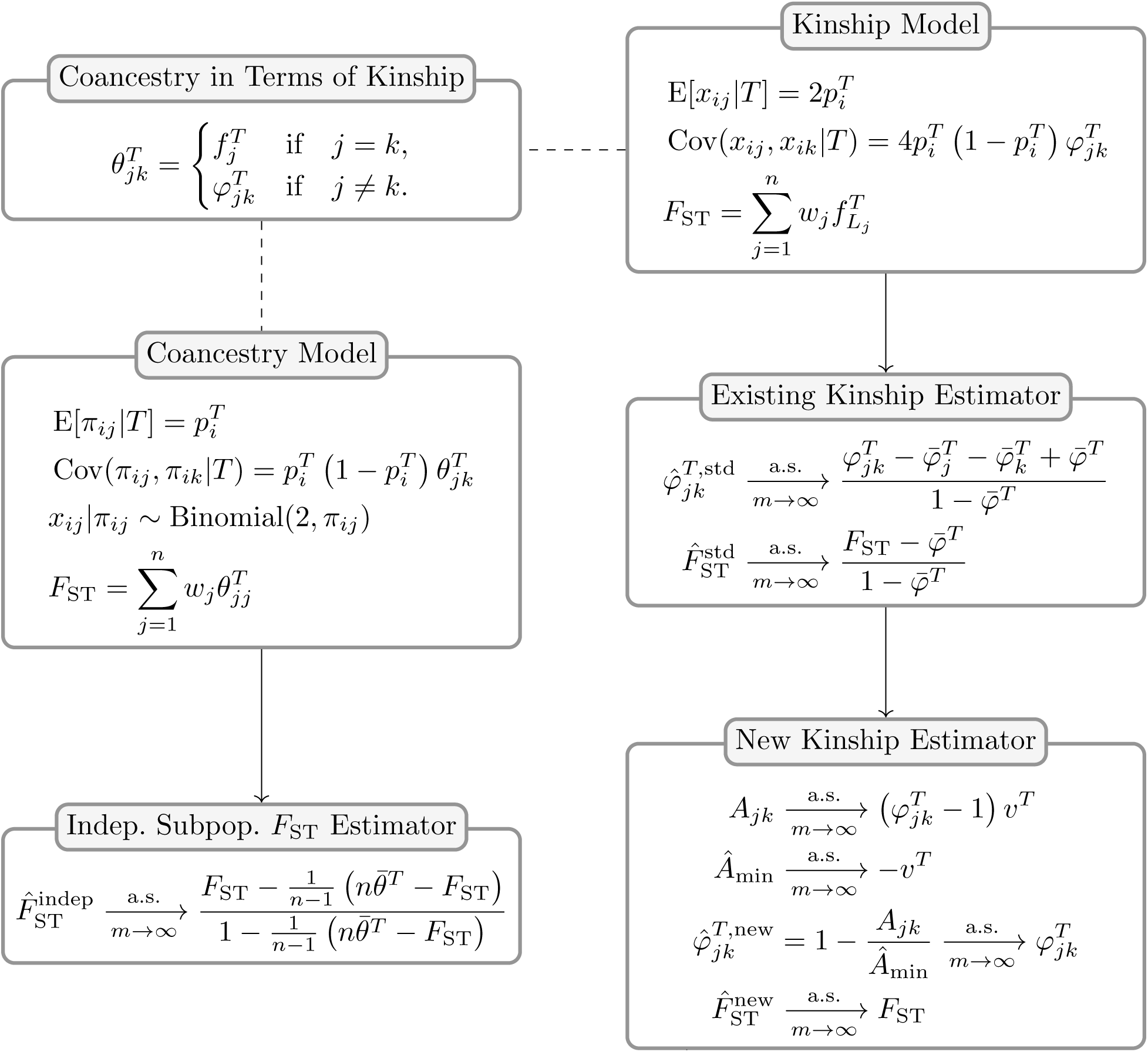
Accuracy of *F*_ST_ and kinship estimators: overview of models and results. Our analysis is based on two parallel models: the “Coancestry Model” for individual-specific allele frequencies (*π*_*ij*_; Section 5 of Part I), and the “Kinship Model” for genotypes (*x*_*ij*_; Section 3.5 of Part I). The “Coancestry in Terms of Kinship” panel connects kinship 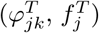 and coancestry 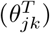 parameters (proven in Section 5.2 of Part I). We use these models to study the accuracy of *F*_ST_ and kinship method-of-moment estimators under arbitrary population structures. The “Indep. Subpop. *F*_ST_ Estimator” panel shows the bias resulting from the misapplication of *F*_ST_ estimators for independent subpopulations 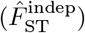 to arbitrary structures (Section 3), as calculated under the coancestry model. The “Existing Kinship Estimator” panel shows the bias in the standard kinship model estimator 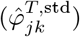 and its resulting plug-in *F*_ST_ estimator (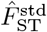; Section 4), as calculated under the kinship model. The “New Kinship Estimator” panel presents a new statistic *A*_*jk*_ that estimates kinship with a uniform bias, which together with a consistent estimator of its minimum value 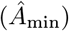 results in our new kinship 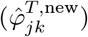 and 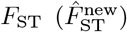 estimators, which are consistent under arbitrary population structure (Section 5). Note that estimation of *F*_ST_ from genotypes requires individuals to be locally outbred and locally unrelated (see Sections 3.2 and 3.3 of Part I).

Here, we study existing *F*_ST_ and kinship method-of-moments estimators in models that allow for arbitrary population structures (see Fig. 1 for an overview of the results). First, in Section 2 we obtain new strong convergence results for a family of ratio estimators that includes the most common *F*_ST_ and kinship estimators. Next, we calculate the convergence values of these *F*_ST_ (Section 3) and kinship (Section 4) estimators under arbitrary population structures, where we find biases that are not present under their original assumptions about structure (panels “Indep. Subpop. *F*_ST_ Estimator” and “Existing Kinship Estimator” in Fig. 1). We characterize the limit of the standard kinship estimator for the first time, identifying complex biases or distortions that have not been described before (related results were independently and concurrently calculated by [53]). In Section 5 we introduce a new approach for kinship and *F*_ST_ estimation for arbitrary population structures, and demonstrate the improved performance using a simple implementation of these estimators (panel “New Kinship Estimator” in Fig. 1). Lastly, in Section 6 we construct an admixture simulation that does not have independent subpopulations to illustrate our theoretical findings through simulation. Elsewhere, we analyze the Human Origins and 1000 Genomes Project datasets with our novel kinship and *F*_ST_ estimation approach, where we demonstrate its coherence with the African Origins model, and illustrate the shortcomings of previous approaches in these complex data [47]. In summary, we identify a new approach for unbiased estimation of *F*_ST_ and kinship, and we provide new estimators that are nearly unbiased.

## 2 Assessing the accuracy of genome-wide estimators

Many *F*_ST_ and kinship coefficient method-of-moments estimators are *ratio estimators*, a general class of estimators that tends to be biased and to have no closed-form expectation [54]. In the *F*_ST_ literature, the expectation of a ratio is frequently approximated with a ratio of expectations [6, 19, 20]. Specifically, ratio estimators are often called “unbiased” if the ratio of expectations is unbiased, even though the ratio estimator itself may be biased [54]. Here we characterize the behavior of two ratio estimator families calculated from genome-wide data, detailing conditions where the previous approximation is justified and providing additional criteria to assess the accuracy of such estimators. These convergence results are the foundation of our analysis of estimators and are applied repeatedly to the various kinship and *F*_ST_ estimators discussed in Sections 3 to 5.

### 2.1 Ratio estimators

The general problem of forming ratio estimators involves random variables *a*_*i*_ and *b*_*i*_ calculated from genotypes at each locus *i*, such that E[*a*_*i*_] = *Ac*_*i*_ and E[*b*_*i*_] = *Bc*_*i*_ and the goal is to estimate 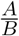. *A* and *B* are constants shared across loci (given by *F*_ST_ or 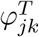), while *c*_*i*_ depends on the ancestral allele frequency 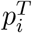 and varies per locus. The problem is that the single-locus estimator 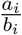 is biased, since 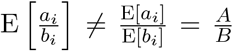, which applies to ratio estimators in general [54]. Below we study two estimator families that combine large numbers of loci to better estimate 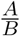.

### 2.2 Convergence

The solution we recommend is the “ratio-of-means” estimator 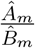, where 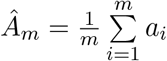, and 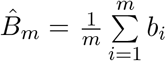, which is common for *F*_ST_ estimators [6, 19, 20, 55]. Note that 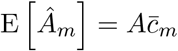 and 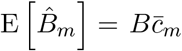, where 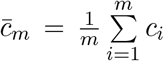. We will assume bounded terms (|*a*_*i*_|, |*b*_*i*_| ≤ *C* for some finite *C*), a convergent 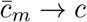, and *Bc* ≠ 0, which are satisfied by common estimators. Given independent loci, we prove almost sure convergence to the desired quantity (Supplementary Information, Section S1.1),

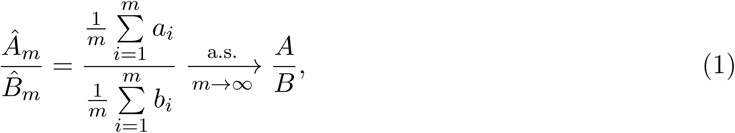

a strong result that implies 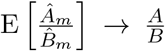, justifying previous work [6, 19, 20]. Moreover, the error between these expectations scales with 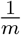 (Supplementary Information, Section S1.2), just as for standard ratio estimators [54]. Although real loci are not independent due to genetic linkage, their dependence is very localized, so this estimator will perform well if the effective number of independent loci is large.

In order to test if a given ratio-of-means estimator converges to its ratio of expectations as in Eq. (1), the following three conditions must be met. (i) The expected values of each term *a*_*i*_, *b*_*i*_ must be calculated and shown to be of the form E[*a*_*i*_] = *Ac*_*i*_ and E[*b*_*i*_] = *Bc*_*i*_ for some *A* and *B* shared by all loci *i* and some *c*_*i*_ that may vary per locus *i* but must be shared by both E[*a*_*i*_], E[*b*_*i*_]. In the estimators we study, *A* and *B* are functions of IBD probabilities such as 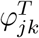 and *F*_ST_, while *c*_*i*_ is a function of 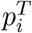 only. (ii) The mean *c*_*i*_ must converge to a non-zero value for infinite loci. (iii) Both |*a*_*i*_|, |*b*_*i*_| ≤ *C* must be bounded for all *i* by some finite *C* (the estimators we study usually have *C* = 1 or *C* = 4). If these conditions are satisfied, then Eq. (1) holds for independent loci and the *A* and *B* found in the first step. See Section 3.2 for an example application of this procedure to an *F*_ST_ estimator.

Another approach is the “mean-of-ratios” estimator 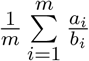, used often to estimate kinship coefficients [18, 24, 27–32] and *F*_ST_ [36]. If each 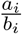 is biased, their average across loci will also be biased, even as *m → ∞*. However, if 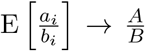 for all loci *i* = 1,…, *m* as the number of individuals *n* → *∞*, and 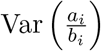 is bounded, then

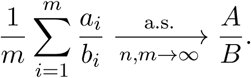

Therefore, mean-of-ratios estimators must satisfy more restrictive conditions than ratio-of-means estimators, as well as large *n* (in addition to the large *m* needed by both estimators), to estimate 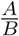 well. We do not provide a procedure to test whether a given mean-of-ratios estimator converges as shown above.

## 3 *F*_ST_ estimation based on the independent subpopulations model

Now that we have detailed how ratio estimators may be evaluated for their accuracy, we turn to existing estimators and assess their accuracy under arbitrary population structures. We study the Weir-Cockerham (WC) [19] and “Hudson” [20] *F*_ST_ estimators, which assume the independent subpopulations model described above. The panel “Indep. Subpop. *F*_ST_ Estimator” in Fig. 1 provides an overview of our results, which we detail in this section.

### 3.1 The *F*_ST_ estimator for independent subpopulations and infinite subpopulation sample sizes

The WC and Hudson method-of-moments estimators have small sample size corrections that remarkably make them consistent as the number of independent loci *m* goes to infinity for finite numbers of individuals. However, these small sample corrections also make the estimators unnecessarily cumbersome for our purposes (see Supplementary Information, Section S2 for complete formulas). In order to illustrate clearly how these estimators behave, both under the independent subpopulations model and for arbitrary structure, here we construct simplified versions that assume infinite sample sizes per subpopulation (see Supplementary Information, Section S2 for details). This simplification corresponds to eliminating statistical sampling, leaving only genetic sampling to analyze [56]. Note that our simplified estimator nevertheless illustrates the general behavior of the WC and Hudson estimators under arbitrary structure, and the results are equivalent to those we would obtain under finite sample sizes of individuals. While the Hudson *F*_ST_ estimator compares two subpopulations [20], we derive a new generalized “HudsonK” estimator for more than two subpopulations in Supplementary Information, Section S2.3.

Under infinite subpopulation sample sizes, the allele frequencies at each locus and every sub-population are known. Let *j* ∈ {1,…, *n*} index subpopulations rather than individuals and *π*_*ij*_ be the allele frequency in subpopulation *j* at locus *i*. We call the *π*_*ij*_ values “individual-specific allele frequencies” (IAF), as has been previously done [49]. In this special case, both WC and HudsonK simplify to the following *F*_ST_ estimator for independent subpopulations (“indep”; derived in Supplementary Information, Section S2):

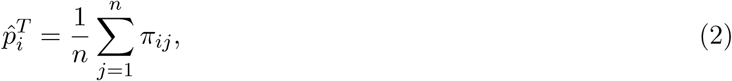

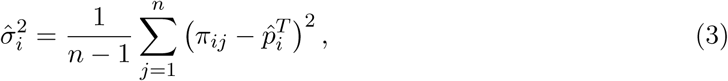

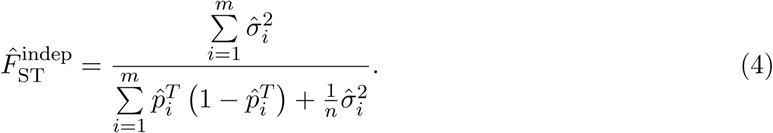

The goal is to estimate 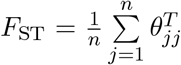, which weighs every subpopulation *j* equally 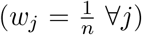, under the coancestry model of Part I, which assumes the following moments for IAFs:

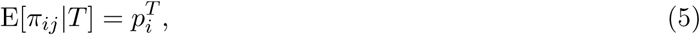

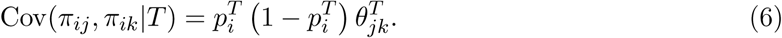

### 3.2 *F*_ST_ estimation under the independent subpopulations model

Under the independent subpopulations model 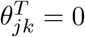 for *j* ≠ *k*, where *T* is the most recent common ancestor (MRCA) population of the set of subpopulations. Note that the estimator in Eq. (4) can be derived directly from Eqs. (5) and (6) and these assumptions using the method of moments (ignoring the existence of previous *F*_ST_ estimators; Supplementary Information, Section S3.1). The expectations of the two recurrent terms in Eq. (4) are

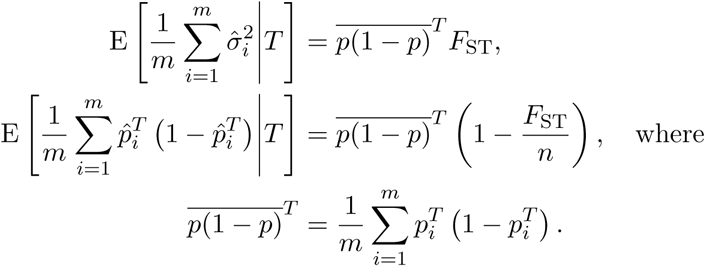

Eliminating 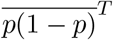 and solving for *F*_ST_ in this system of equations recovers the estimator in Eq. (4).

Before applying the convergence result in Eq. (1), we test that the three conditions listed in Section 2 are met. Condition (i): The locus *i* terms are 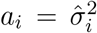 and 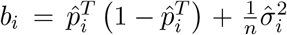, which satisfy E[*a*_*i*_] = *Ac*_*i*_ and E[*b*_*i*_] = *Bc*_*i*_ with *A* = *F*_ST_, *B* = 1, and 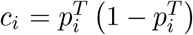. Condition (ii): 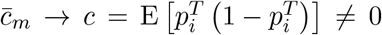 over the 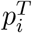 distribution across loci. Condition (iii): Since 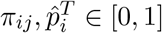, then 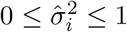 and 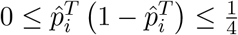, and since *n* ≥ 2, *C* = 1 bounds both |*a*_*i*_| and |*b*_*i*_|. Therefore, for independent loci,

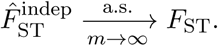

### 3.3 *F*_ST_ estimation under arbitrary coancestry

Now we consider applying the independent subpopulations *F*_ST_ estimator to dependent subpopulations. The key difference is that now 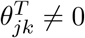 for every (*j, k*) will be assumed in our coancestry model in Eqs. (5) and (6), and now *T* may be either the MRCA population of all individuals or a more ancestral population. In this general setting, (*j, k*) may index either subpopulations or individuals. The two terms of 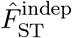 now satisfy

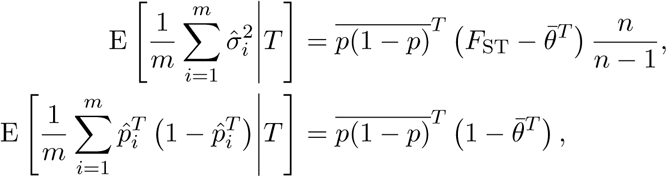

where 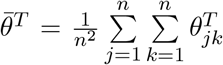 is the mean coancestry with uniform weights. There are two equations but three unknowns: *F*_ST_, and 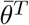, and 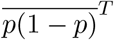. The independent subpopulations model satisfies 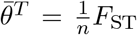, which allows for the consistent estimation of *F*_ST_. Therefore, the new unknown 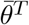 precludes consistent *F*_ST_ estimation without additional assumptions.

The *F*_ST_ estimator for independent subpopulations converges more generally to

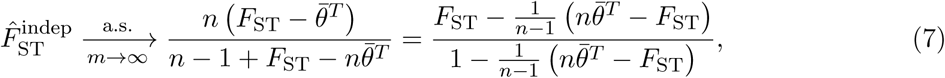

(the conclusion of panel “Indep. Subpop. *F*_ST_ Estimator” in Fig. 1), where it should be noted that

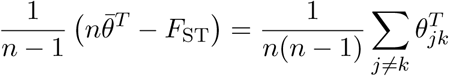

is the average of all between-individual coancestry coefficients, a term that appears in a related result for subpopulations [6]. Therefore, under arbitrary structure the independent subpopulations estimator’s bias is due to the coancestry between individuals (or subpopulations in the traditional setting). While the limit in Eq. (7) appears to vary depending on the choice of *T*, it is in fact a constant with respect to *T* (proof in Supplementary Information, Section S4.1).

Since 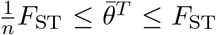 (Supplementary Information, Section S5), this estimator has a downward bias in the general setting: it is asymptotically unbiased 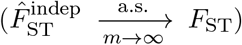 only when 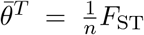, while bias is maximal when 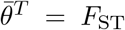, where 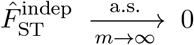. For example, if 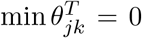 so the MRCA population *T* is fixed, but *n* is large and 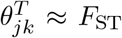 for most pairs of individuals, then 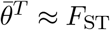 as well, and 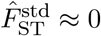. Therefore, the magnitude of the bias of 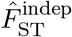 is unknown if 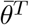 is unknown, and small 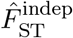 may arise even if *F*_ST_ is very large.

### 3.4 Coancestry estimation as a method of moments

Since the generalized *F*_ST_ is given by coancestry coefficients 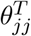 (Eq. (13) of Part I), a new *F*_ST_ estimator could be derived from estimates of 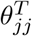. Here we attempt to define a method-of-moments estimator for 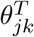, and find an underdetermined estimation problem, just as for *F*_ST_.

Given IAFs and Eqs. (5) and (6), the first and second moments that average across loci are

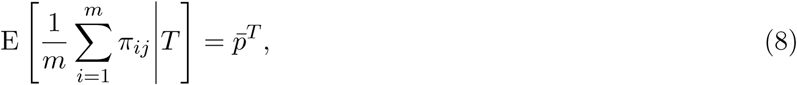

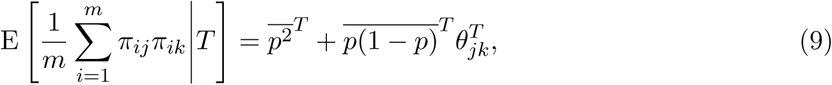

where 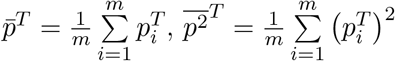, and 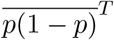 is as before.

Suppose first that only 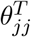 are of interest. There are *n* estimators given by Eq. (9) with *j* = *k*, each corresponding to an unknown 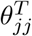. However, all these estimators share two nuisance parameters: 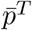 and 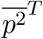. While 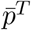 can be estimated from Eq. (8), there are no more equations left to estimate 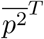, so this system is underdetermined. The estimation problem remains underdetermined if all 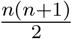 estimators in Eq. (9) are considered rather than only the *j* = *k* cases. Therefore, we cannot estimate coancestry coefficients consistently using only the first two moments without additional assumptions.

## 4 Characterizing a kinship estimator and its relationship to *F*_ST_

Given the biases we see for 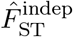 under arbitrary structures in Section 3.3, we now turn to the generalized definition of *F*_ST_ and pursue an estimate of it. Recall from Eq. (3) of Part I that our generalized *F*_ST_ is defined in terms of inbreeding coefficients, which are a special case of the kinship coefficient:

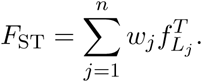

Therefore, we will first consider estimates of kinship and inbreeding in this section. Note also that estimating kinship is important for GWAS approaches that control for population structure [18, 21–32, 57, 58]. Lastly, kinship coefficients determine the bias of 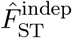 in Eq. (7) (since coancestry and kinship coefficients are closely related: see panel “Coancestry in Terms of Kinship” in Fig. 1).

In this section, we focus on a standard kinship method-of-moments estimator and calculate its limit for the first time (panel “Existing Kinship Estimator” in Fig. 1). We study estimators that use genotypes or IAFs, and construct *F*_ST_ estimators from their kinship estimates. We find biases comparable to those of 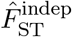 (Section 3), and define unbiased *F*_ST_ estimators that require knowing the mean kinship or coancestry, or its proportion relative to *F*_ST_. The results of this section directly motivate and help construct our new kinship and *F*_ST_ estimation approach in Section 5.

### 4.1 Characterization of the standard kinship estimator

Here we analyze a standard kinship estimator that is frequently used [18, 24, 27–33]. We generalize this estimator to use weights in estimating the ancestral allele frequencies, and we write it as a ratio-of-means estimator due to the favorable theoretical properties of this format as detailed in Section 2:

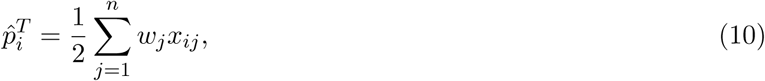

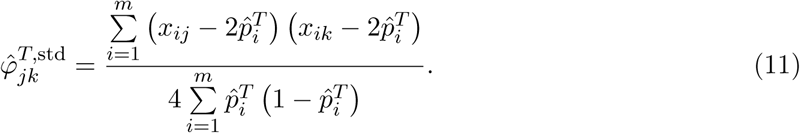

The estimator in Eq. (11) resembles the sample covariance estimator applied to genotypes, but centers by locus *i* rather than by individuals *j* and *k*, and normalizes using estimates of 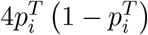. We derive Eq. (11) directly using the method of moments in Supplementary Information, Section S3.2. The weights in Eq. (10) must satisfy *w*_*j*_ > 0 and 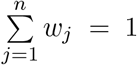, so 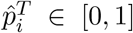 and 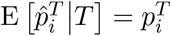.

Utilizing the following moments for genotypes (from the kinship model of Part I),

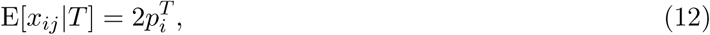

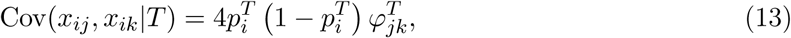

we find that Eq. (11) converges to

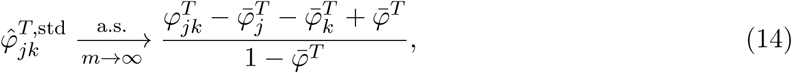

where 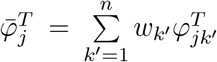 and 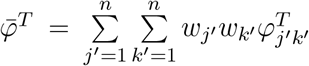. (This is the conclusion of panel “Existing Kinship Estimator” in Fig. 1; see Supplementary Information, Section S6 for intermediate calculations that lead to Eq. (14).) Therefore, the bias of 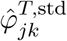 varies per *j* and *k*. Analogous distortions have been observed for sample covariances of genotypes [59] and were found in concurrent independent work [53]. The limit of 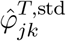 in Eq. (14) is constant with respect to *T* (proof in Supplementary Information, Section S4.2). Similarly, inbreeding coefficient estimates derived from Eq. (11) converge to

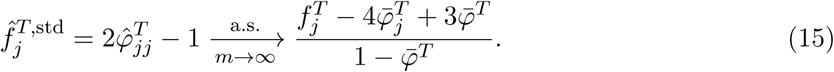

The difference between the bias of 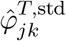 for *j* ≠ *k* in Eq. (14) and 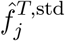 in Eq. (15) is visible in the kinship estimates of Fig. 5C (the difference causes a discontinuity between the diagonal and off-diagonal values). The limits of the ratio-of-means versions of two more 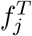 estimators [29] are, if 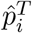 uses Eq. (10),

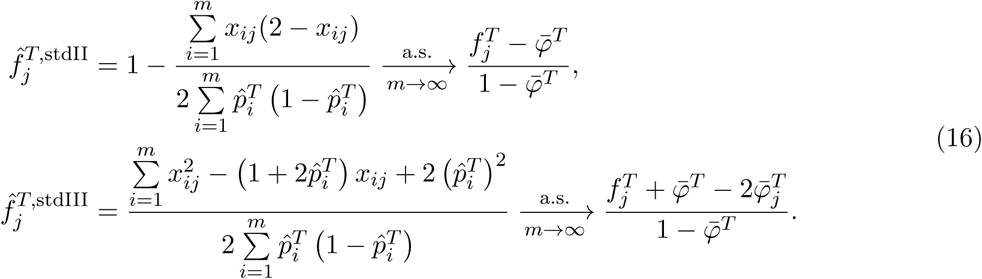

The estimators in Eqs. (11) and (16) are unbiased when 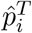 is replaced by 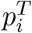 [18, 29, 33], and are consistent when 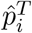 is consistent [48]. Surprisingly, 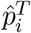 in Eq. (10) is not consistent (it does not converge almost surely) for arbitrary population structures, which is at the root of the bias in Eqs. (14) to (16). In particular, although 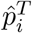 is unbiased, its variance (see Supplementary Information, Section S6),

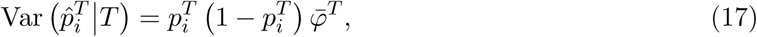

may be asymptotically non-zero as *n* → *∞*, since 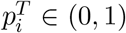 is fixed and 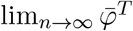 may take on any value in [0,1] for arbitrary population structures. Further, 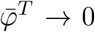 as *n* → *∞* if and only if 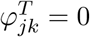 for almost all pairs of individuals (*j, k*). These observations hold for any weights such that *w*_*j*_ > 0, 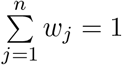. An important consequence is that the plug-in estimate of 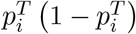 is biased (Supplementary Information, Section S6),

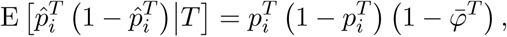

which is present in all estimators we have studied.

### 4.2 Estimation of coancestry coefficients from IAFs

Here we form a coancestry coefficient estimator analogous to Eq. (11) but using IAFs. Assuming the moments in Eqs. (5) and (6), this estimator and its limit are

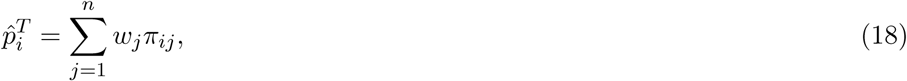

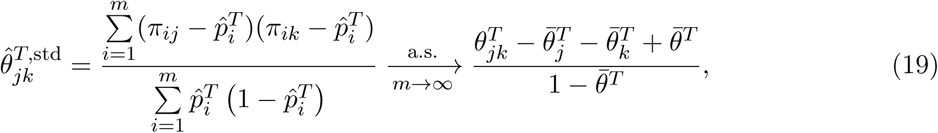

where 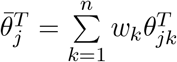 and 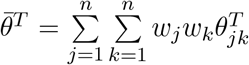 are analogous to 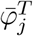 and 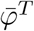. Eq. (18) generalizes Eq. (2) for arbitrary weights. Thus, use of IAFs does not ameliorate the estimation problems we have identified for genotypes. Like Eq. (17), 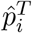 in Eq. (18) is not consistent because 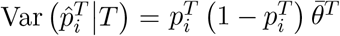 may not converge to zero for arbitrary population structures, which causes the bias observed in Eq. (19).

### 4.3 *F*_ST_ estimator based on the standard kinship estimator

Since the generalized *F*_ST_ is defined as a mean inbreeding coefficient (Eq. (3) of Part I), here we study the *F*_ST_ estimator constructed as 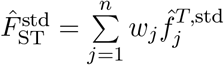 where 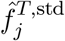 is the inbreeding estimator derived from the standard kinship estimator. Although 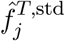 is biased, we nevertheless plug it into our definition of *F*_ST_ so that we may study how bias manifests. Note that we do not recommend utilizing this *F*_ST_ estimator in practice, but we find these results informative for identifying how to proceed in deriving new estimators (Section 5).

Remarkably, the three 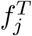 estimators in Eqs. (15) and (16) give exactly the same plug-in 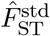 if the weights in *F*_ST_ and 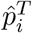 in Eq. (10) match, namely

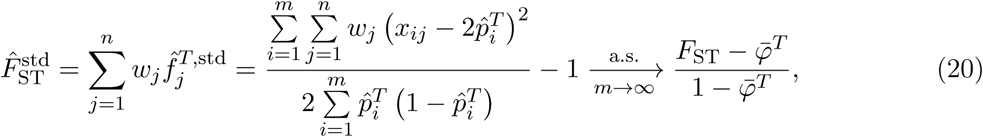

where the limit assumes locally-outbred individuals so 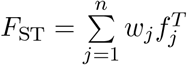. The analogous *F*_ST_ estimator for IAFs and its limit are

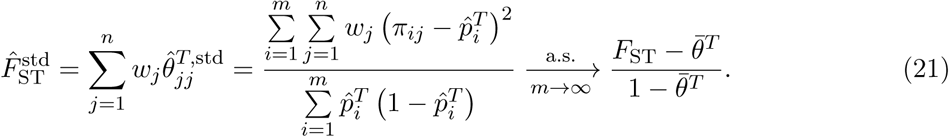

The estimators in Eqs. (20) and (21) for individuals and their limits resemble those of classical *F*_ST_ estimators for populations of the form 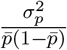 [6, 7].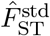 in Eq. (21) for subpopulations *j* with uniform weight and one locus is also *G*_ST_ for two alleles [60]. Compared to 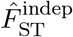 in Eq. (4), 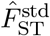 in Eq. (21) admits arbitrary weights and, by forgoing bias correction under the independent subpopulations model, is a simpler target of study.

Like 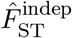 in Eq. (4), 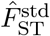 in Eqs. (20) and (21) are downwardly biased since 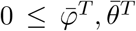. 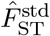 in Eq. (21) may converge arbitrarily close to zero since 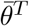 can be arbitrarily close to *F*_ST_ (Supplementary Information, Section S5). Moreover, although 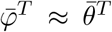 for large *n* (see panel “Coancestry in Terms of Kinship” in Fig. 1), in extreme cases 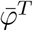 can exceed *F*_ST_ under the coancestry model (where 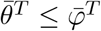) and also under extreme local kinship, where 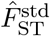 in Eq. (20) converges to a negative value.

### 4.4 Adjusted consistent oracle *F*_ST_ estimators and the “bias coefficient”

Here we explore two adjustments to 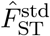 from IAFs in Eq. (21) that rely on having minimal additional information needed to correct its bias. If 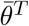 is known, the bias in Eq. (21) can be reversed, yielding the consistent estimator

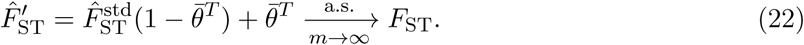

Consistent estimates are also possible if a scaled version of 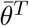 is known, namely

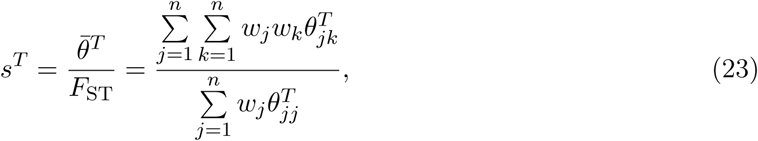

which we call the “bias coefficient” and which has interesting properties. The bias coefficient quantifies the departure from the independent subpopulations model by comparing the mean coancestry 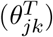 to the mean inbreeding coefficient 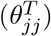, and given *F*_ST_ > 0 satisfies 0 < *s*^*T*^ ≤ 1 (Supplementary Information, Section S5). The limit in Eq. (21) in terms of *s*^*T*^ is

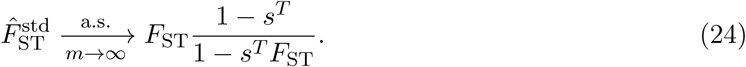

Treating the limit as equality and solving for *F*_ST_ yields the following consistent estimator:

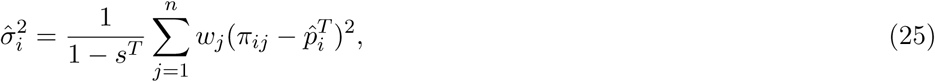

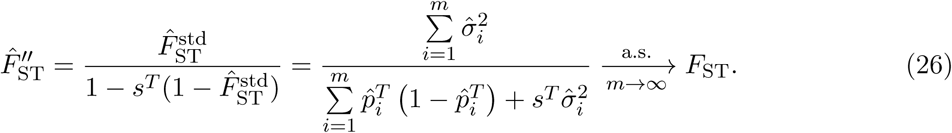

Note that 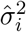 and 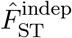 from Eqs. (3) and (4) are the special case of Eqs. (25) and (26) for uniform weights and 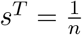; hence, 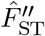 generalizes 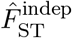.

Lastly, using either Eq. (21) or Eq. (24), the relative error of 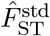 converges to

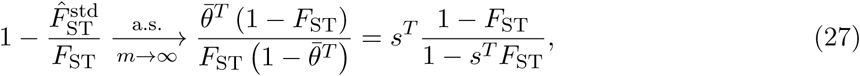

which is approximated by *s*^*T*^ if *F*_ST_ « 1, hence the name “bias coefficient”. Note *s*^*T*^ varies depending on the choice of *T*, which is necessary since *F*_ST_ (and hence the relative bias of 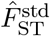 from *F*_ST_) depends on the choice of *T*.

## 5 A new approach for kinship and *F*_ST_ estimation

Here, we propose a new estimation approach for kinship coefficients that has properties favorable for obtaining nearly unbiased estimates (panel “New Kinship Estimator” in Fig. 1). These new kinship estimates yield an improved *F*_ST_ estimator. We present the general approach and implement a simple version of one key estimator that results in the complete proof-of-principle estimator that is evaluated in Section 6 and applied to human data in [47].

### 5.1 General approach

In this subsection we develop our new estimator in two steps. First, we compute a new statistic *A*_*jk*_ that is proportional in the limit of infinite loci to 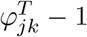 times a nuisance factor *v*^*T*^. Second, we estimate and remove *v*^*T*^ to yield the proposed estimator 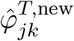. 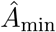—an estimator of the limit of the minimum *A*_*jk*_—yields *v*^*T*^ if the least related pair of individuals in the data has 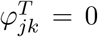, which sets *T* to the MRCA population of all the individuals in the data. The new kinship estimator immediately results in new inbreeding 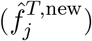 and *F*_ST_ 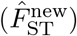 estimators. This general approach leaves the implementation of 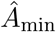; the simple implementation applied in this work is described in Section 5.2, but our method can be readily improved by substituting in a better 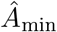 in the future.

Applying the method of moments to Eqs. (12) and (13), we derive the following statistic (see Supplementary Information, Section S7), whose expectation is proportional to 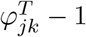:

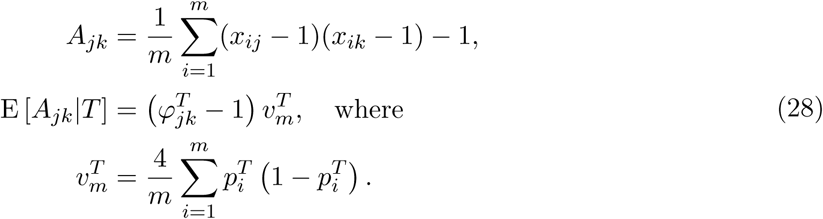

Compared to the standard kinship estimator in Eq. (14), which has a complex asymptotic bias determined by *n* parameters (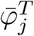 for each *j* ∈ {1,…, *n*}), the *A*_*jk*_ statistics estimate kinship with a bias controlled by the sole unknown parameter 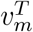 shared by all pairs of individuals. The key to estimating 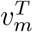 is to notice that if 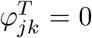 then 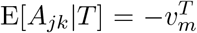. Thus, assuming 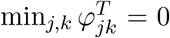, which sets *T* to the MRCA population, then the minimum *A*_*jk*_ yields the nuisance parameter. However, we recommend using a more stable estimate than the minimum *A*_*jk*_ to unbias all *A*_*jk*_, such as the estimator presented in Section 5.2.

In general, suppose 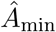 is a consistent estimator of the limit of the minimum *E*[*A*_*jk*_|*T*], or equivalently,

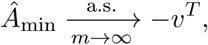

along with the assumption that 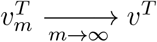 for some *v*^*T*^ ≠ 0. Our new kinship estimator follows directly from replacing 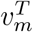 with 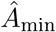 and solving for 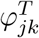 in Eq. (28), which results in a consistent kinship estimator (given the convergence proof of Section 2):

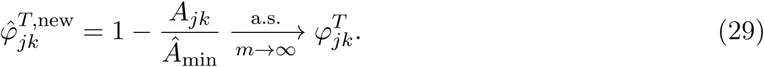

The resulting new inbreeding coefficient estimator is

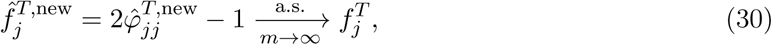

and the new *F*_ST_ estimator is

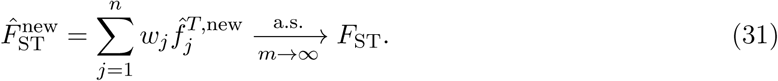

Thus, only the implementation of 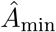 is left unspecified from this general estimation approach of kinship and *F*_ST_. The implementation of 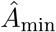 used in the analyses in this work is given in the next subsection.

The *A*_*jk*_ statistic defined above is closely related to the mean “identity by state” estimator [18] and to another recently-described kinship estimator [53, 61]. However, only our 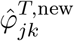 in Eq. (29)—scaling *A*_*jk*_ using 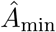—results in consistent kinship estimation under arbitrary population structures.

### 5.2 Proof-of-principle kinship estimator using subpopulation labels

To showcase the potential of the new estimators, we implement a simple proof-of-principle version of 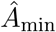 needed for our new kinship estimator (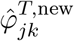 in Eq. (29)). This *Â*_min_ relies on an appropriate partition of the *n* individuals into *K* subpopulations (denoted *S*_*u*_ for *u* ∈ {1,…, *K*}), where the only requirement is that the kinship coefficients between pairs of individuals across the two most unrelated subpopulations is zero, as detailed below. Note that, unlike the the independent subpopulations model of Section 3, these *K* subpopulations need not be independent nor unstructured. The desired estimator *Â*_min_ is the minimum average *A*_*jk*_ over all subpopulation pairs:

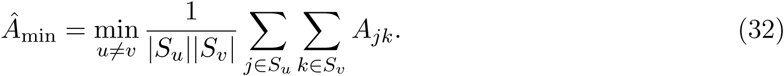

This *Â*_min_ consistently estimates the limit of the minimum *A*_*jk*_ if 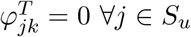, ∀*k* ∈ *S*_*v*_ for the least related pair of subpopulations *S*_*u*_, *S*_*v*_.

This estimator should work well for individuals truly divided into subpopulations, but may be biased for a poor choice of subpopulations, in particular if the minimum mean 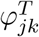 between subpopulations is far greater than zero. For this reason, inspection of the kinship estimates is required and careful construction of appropriate subpopulations may be needed. See our analysis of human data for detailed examples [47]. Future work could focus on a more general 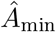 that circumvents the need for subpopulations of our proof-of-principle estimator.

## 6 Simulations evaluating *F*_ST_ and kinship estimators

### 6.1 Overview of simulations

We simulate genotypes from two models to illustrate our results when the true population structure parameters are known. The first simulation satisfies the independent subpopulations model that existing *F*_ST_ estimators assume. The second simulation is from an admixture model with no independent subpopulations and pervasive kinship designed to induce large downward biases in existing kinship and *F*_ST_ estimators (Fig. 2). This admixture scenario resembles the population structure we estimated for Hispanics in the 1000 Genomes Project [47]: compare the simulated kinship matrix (Fig. 2B) and admixture proportions (Fig. 3C) to our estimates on the real data [47]. Both simulations have *n* = 1000 individuals, *m* = 300, 000 loci, and *K* = 10 subpopulations or intermediate subpopulations. These simulations have *F*_ST_ = 0.1, comparable to previous estimates between human populations (in 1000 Genomes, the estimated *F*_ST_ between CEU (European-Americans) and CHB (Chinese) is 0.106, between CEU and YRI (Yoruba from Nigeria) it is 0.139, and between CHB and YRI it is 0.161 [20]).

**Figure 2:**
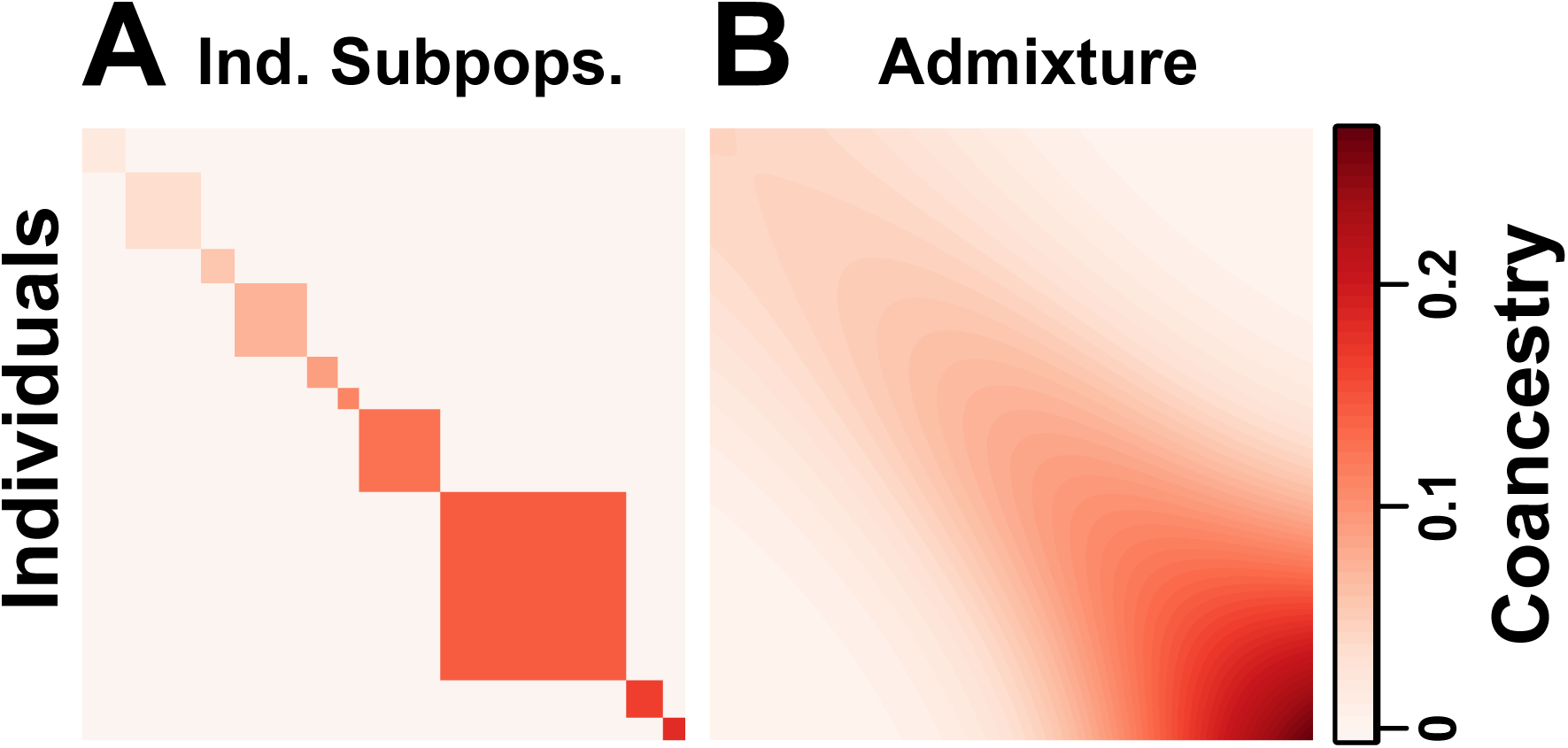
Coancestry matrices of simulations. Both panels have *n* = 1000 individuals along both axes, *K* = 10 subpopulations (final or intermediate), and *F*_ST_ = 0.1. Color corresponds to 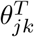 between individuals *j* and *k* (equal to 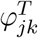 off-diagonal, 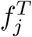 along the diagonal). A) The independent subpopulations model has 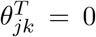 between subpopulations, and varying 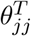 per subpopulation, resulting in a block-diagonal coancestry matrix. B) Our admixture scenario models a 1D geography with extensive admixture and intermediate subpopulation differentiation that increases with distance, resulting in a smooth coancesty matrix with no independent subpopulations (no 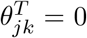 between blocks). Individuals are ordered along each axis by geographical position.

**Figure 3:**
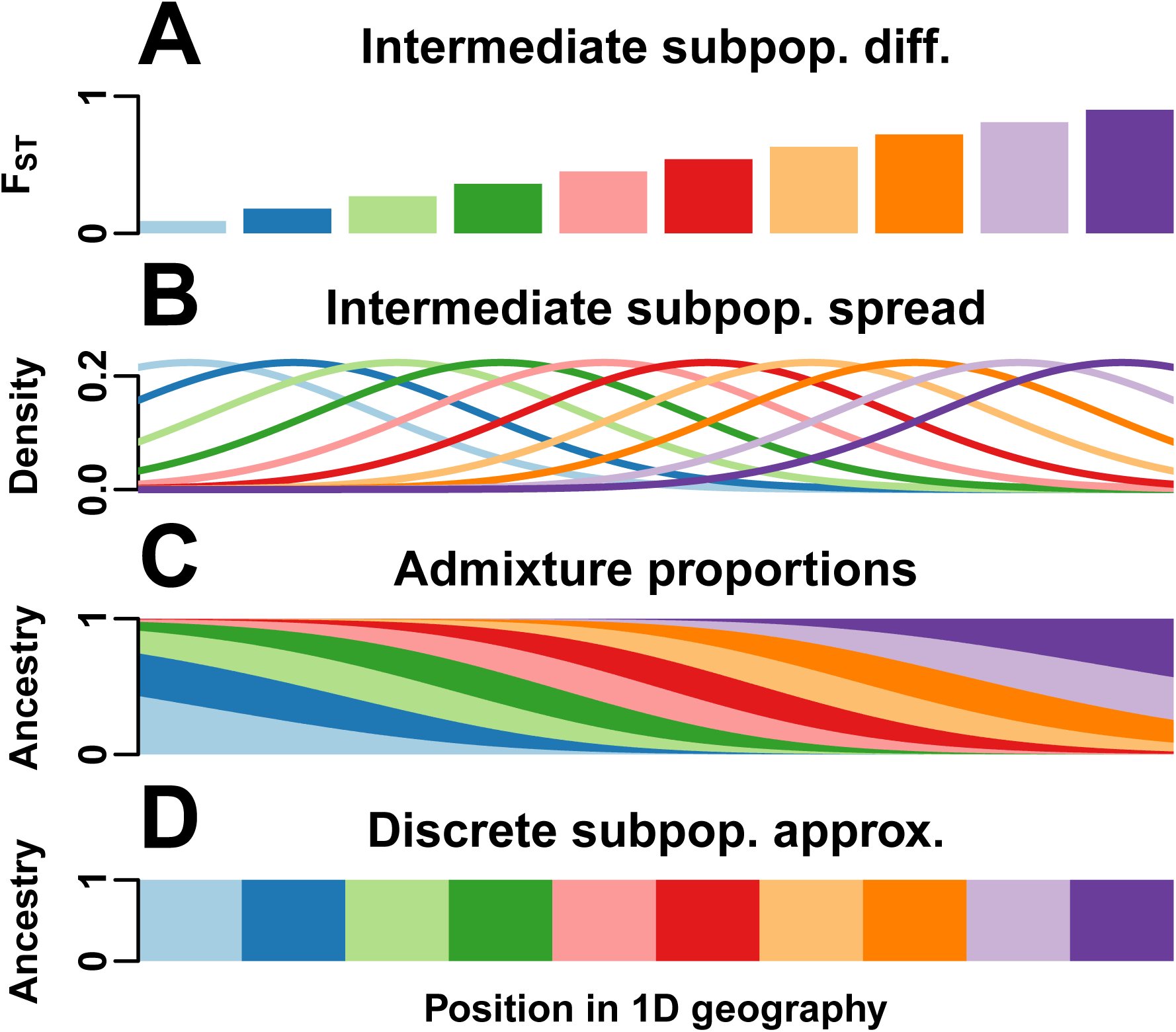
1D admixture scenario. We model a 1D geography population that departs strongly from the independent subpopulations model. A) *K* = 10 intermediate subpopulations, evenly spaced on a line, evolved independently in the past with *F*_ST_ increasing with distance, which models a sequence of increasing founder effects (from left to right) to mimic the global human population. B) Once differentiated, individuals in these intermediate subpopulations spread by random walk modeled by Normal densities. C) *n* = 1000 individuals, sampled evenly in the same geographical range, are admixed proportionally to the previous Normal densities. Thus, each individual draws most of its alleles from the closest intermediate subpopulation, and draws the fewest alleles from the most distant populations. Long-distance random walks of intermediate subpopulation individuals results in kinship for admixed individuals that decays smoothly with distance in Fig. 2B. D) For *F*_ST_ estimators that require a partition of individuals into subpopulations, individuals are clustered by geographical position (*K* = 10).

The independent subpopulations simulation satisfies the HudsonK and BayeScan estimator assumptions: each independent subpopulation *S*_*u*_ has a different *F*_ST_ value of 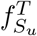 relative to the MRCA population *T* (Fig. 2A). Ancestral allele frequencies 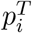 are drawn uniformly in [0.01, 0.5]. Allele frequencies 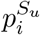 for *S*_*u*_ and locus *i* are drawn independently from the Balding-Nichols (BN) distribution [5] with parameters 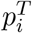 and 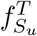. Every individual *j* in subpopulation *S*_*u*_ draws alleles randomly with probability 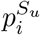. Subpopulation sample sizes are drawn randomly (Supplementary Information, Section S8).

The admixture simulation corresponds to a “BN-PSD” model [8, 24, 31, 48, 62], which we analyzed in Section 6 of Part I and has a demographic model illustrated in Fig. 4 of Part I. The intermediate subpopulations are independent subpopulations that draw 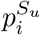 from the BN model, then each individual *j* constructs its allele frequencies as 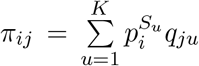, which is a weighted average of 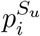 with the admixture proportions *q*_*ju*_ of *j* and *u* as weights (which satisfy 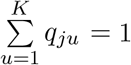, as in the Pritchard-Stephens-Donnelly [PSD] admixture model [50–52]). We constructed *q*_*ju*_ that model admixture resulting from spread by random walk of the intermediate subpopulations along a one-dimensional geography, as follows. Intermediate subpopulations *S*_*u*_ are placed on a line with differentiation 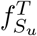 that grows with distance, which corresponds to a serial founder effect (Fig. 3A). Upon differentiation, individuals in each *S*_*u*_ spread by random walk, a process modeled by Normal densities (Fig. 3B). Admixed individuals derive their ancestry proportional to these Normal densities, resulting in a genetic structure governed by geography (Fig. 3C, Fig. 2B) and departing strongly from the independent subpopulations model (Fig. 3D). The amount of spread— which sets the mean kinship across all individuals—was chosen to give a bias coefficient of 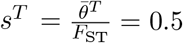, which by Eq. (27) results in a large downward bias for 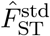 (in contrast, the independent subpopulations simulation has *s*^*T*^ = 0.1). The true 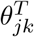 and *F*_ST_ parameters of this simulation are given by the 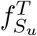 values of the intermediate subpopulations and the admixture coefficients *q*_*ju*_ of the individuals via Eq. (17) of Part I. See Supplementary Information, Section S8 for additional details regarding these simulations.

**Figure 4:**
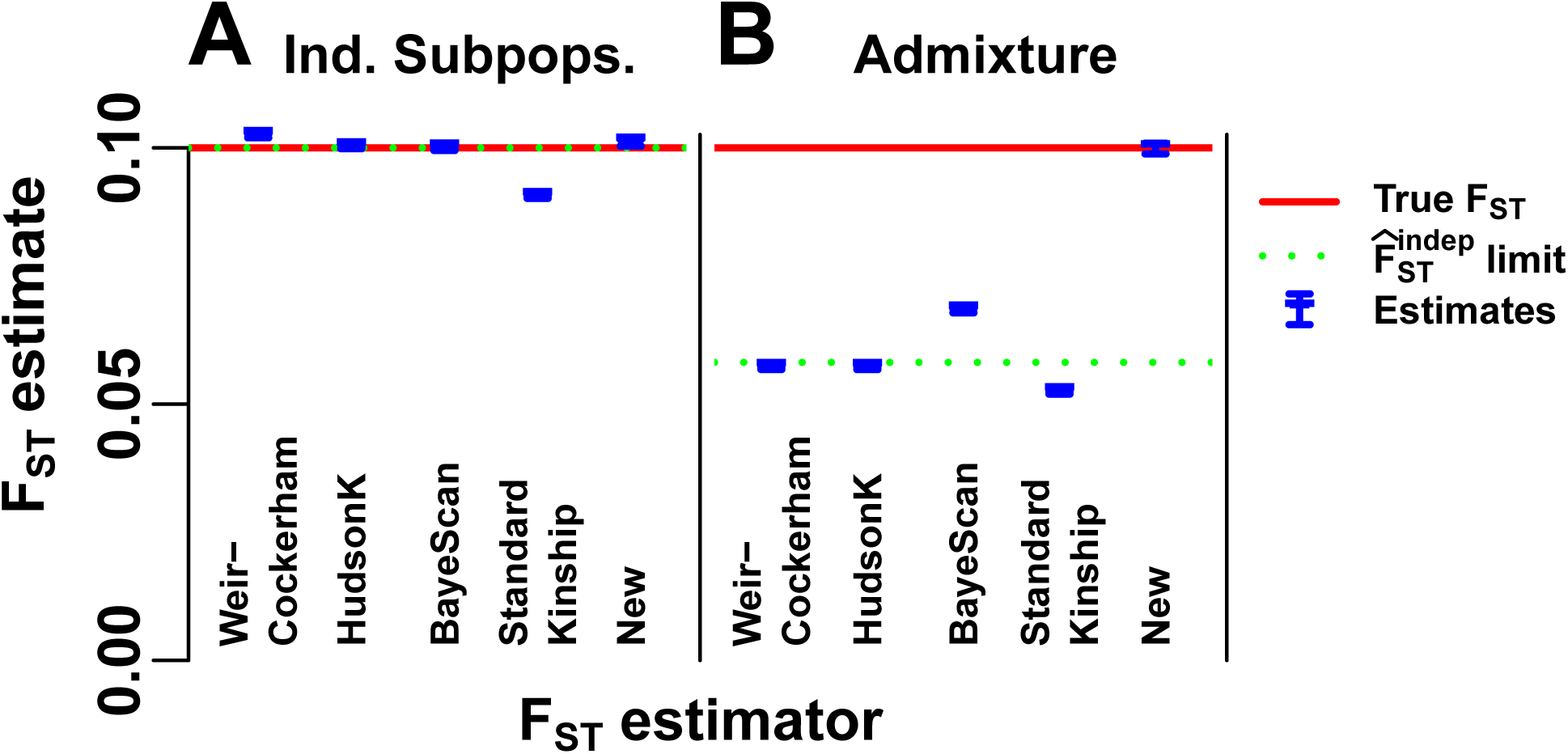
Evaluation of *F*_ST_ estimators. The WC, HudsonK, BayeScan, 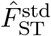 in Eq. (20) derived from the standard kinship estimator, and our new *F*_ST_ estimator in Eqs. (29) and (32), are evaluated on simulated genotypes from our two models (Fig. 2). A) The independent subpopulations model assumed by the HudsonK and BayeScan *F*_ST_ estimators. All but standard kinship 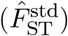 have zero or small biases. B) Our admixture scenario, which has no independent subpopulations, was constructed so 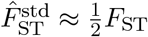. Only our new estimates are accurate. The rest of these estimators have large biases that result from treating kinship as zero between every subpopulations imposed by geographic clustering. The estimator limit in Eq. (7) (green dotted line) overlaps the true *F*_ST_ (red line) in (A) but not (B). Estimates (blue) include 95% prediction intervals (often too narrow to see) from 39 independently-simulated genotype matrices for each model (Supplementary Information, Section S9).

### 6.2 Evaluation of *F*_ST_ estimators

Our admixture simulation illustrates the large biases that can arise if *F*_ST_ estimators for independent subpopulations (WC, HudsonK and BayeScan) are misapplied to arbitrary population structures to estimate the generalized *F*_ST_, and demonstrate the higher accuracy of our new *F*_ST_ estimator 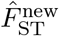 given by the combination of Eqs. (31) and (32). BayeScan was used to estimate the per-subpopulation *F*_ST_ across loci assuming no selection, and the global *F*_ST_ was given by the mean *F*_ST_ across subpopulations.

First, we test these estimators in our independent subpopulations simulation. Both the HudsonK (Supplementary Information, Section S2.3) and BayeScan *F*_ST_ estimators are consistent in this simulation, since their assumptions are satisfied (Fig. 4A). The WC estimator assumes that 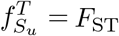 for all subpopulations *S*_*u*_, which does not hold; nevertheless, WC has only a small bias (Fig. 4A). For comparison, we show the standard kinship-based 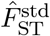 in Eq. (20) (weights from Supplementary Information, Section S8), which does not have corrections that would make it consistent under the independent subpopulations model. Since the number of subpopulations *K* is large, 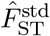 has a small relative bias of about 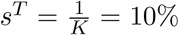 (Fig. 4A); greater bias is expected for smaller *K*. Our new *F*_ST_ estimator has a very small bias in this simulation resulting from estimating the minimum kinship from the smallest kinship between subpopulations (see Eq. (32)) rather than their average as HudsonK does implicitly (Fig. 4A).

Next we test these estimators in our admixture simulation. To apply the *F*_ST_ estimators that require subpopulations to the admixture model, individuals are clustered into subpopulations by their geographical position (Fig. 3D). We find that estimates of WC, HudsonK, and BayeScan are smaller than the true *F*_ST_ by nearly half, as predicted by the limit of 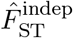 in Eq. (7) (Fig. 4B). By construction, 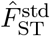 also has a large relative bias of about *s*^*T*^ = 50%; remarkably, the WC, HudsonK, and BayeScan estimators suffer from comparable biases. Thus, the corrections for independent subpopulations present in the WC and HudsonK estimators, or the Bayesian likelihood modeling of BayeScan, are insufficient for accurate estimation of the generalized *F*_ST_ in this admixture scenario. Only our new *F*_ST_ estimator achieves practically unbiased estimates in the admixture simulation (Fig. 4B).

### 6.3 Evaluation of kinship estimators

Our admixture simulation illustrates the distortions of the standard kinship estimator 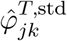 in Eq. (11) and demonstrates the improved accuracy of our new kinship estimator 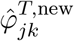 given by the combination of Eqs. (29) and (32). The limit of the standard estimator 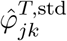 in Eq.(11) has a uniform bias if 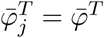 for all individuals *j*. For that reason, our admixture simulation has varying differentiation 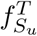 per intermediate subpopulation *S*_*u*_ (Fig. 3A), which causes large differences in 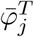 per individual *j* and therefore large distortions in 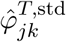.

Our new kinship estimator (Fig. 5B) recovers the true kinship matrix of this complex population structure (Fig. 5A), with an RMSE of 2.83% relative to the mean 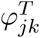. In contrast, estimates using the standard estimator have a large overall downward bias (Fig. 5C), resulting in an RMSE of 115.72% from the true 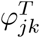 relative to the mean 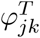. Additionally, estimates from 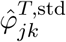 are very distorted, with an abundance of 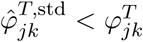 cases—some of which are negative estimates (blue in Fig. 5C)—but remarkably also cases with 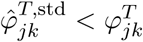(top left corner of Fig. 5C).

**Figure 5:**
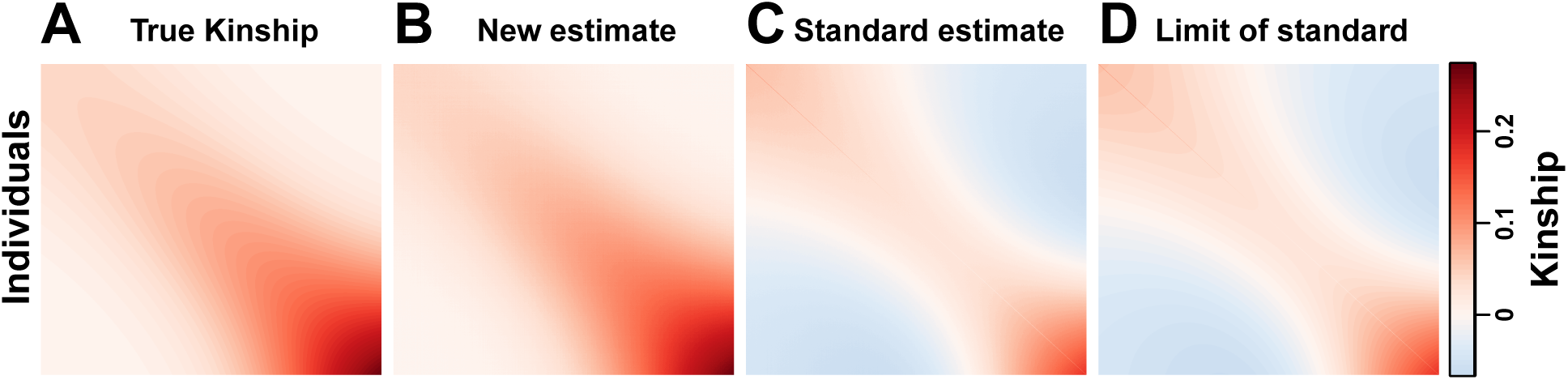
Evaluation of kinship estimators. Bias for the standard kinship coefficient estimator is illustrated in our admixture simulation and contrasted to the nearly unbiased estimates of our new estimator. Plots show *n* = 1000 individuals along both axes, and color corresponds to 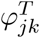 between individuals *j* ≠ *k* and to 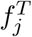 along the diagonal (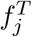 is in the same scale as 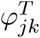 for *j* ≠ *k*; plotting 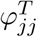, which have a minimum value of 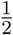, would result in a discontinuity in this figure). A) True kinship matrix. B) Estimated kinship using our new estimator in Eqs. (29) and (32) from simulated genotypes recovers the true kinship matrix with high accuracy. C) Standard kinship estimates 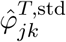 given by Eq. (11) from simulated genotypes are downwardly biased on average and distorted by pair-specific amounts. D) Theoretical limit of 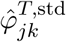 in Eq. (14) as the number of independent loci goes to infinity demonstrates the accuracy of our bias predictions under the kinship model.

Now we compare the convergence of the ratio-of-means and mean-of-ratios versions of the standard kinship estimator to their biased limit we calculated in Eq. (14) (Fig. 5D). The ratio-of-means estimate 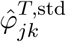 (Fig. 5C) has an RMSE of 2.14% from its limit relative to the mean 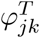. In contrast, the mean-of-ratios estimates that are prevalent in the literature have a greater RMSE of 10.77% from the same limit in Eq. (14). Thus, as expected from our theoretical results in Section 2, the ratio-of-means estimate is much closer to the desired limit than the mean-of-ratio estimate. The distortions are similar for the estimator that uses IAFs in Eq. (19), with reduced RMSEs from its limit of 0.32% and 8.82% for the ratio-of-means and mean-of-ratios estimates, respectively.

### 6.4 Evaluation of oracle adjusted *F*_ST_ estimators

Here we verify additional calculations for the bias of the standard kinship-based estimator 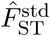 and the unbiased adjusted “oracle” *F*_ST_ estimators that require the true mean kinship 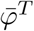 or the bias coefficient *s*^*T*^ to be known. Note that 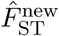 in Eq. (31) is related but not identical to these oracle estimators. We tested both IAF (Fig. 6A) and genotype (Fig. 6B) versions of these estimators. The unadjusted 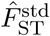 in Eq. (21) is severely biased (blue in Fig. 6) by construction, and matches the calculated limit for IAFs and genotypes (green lines in Fig. 6, which are close because 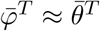). In contrast, the two consistent adjusted estimators 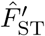 and 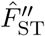 in Eqs. (22) and (26) estimate *F*_ST_ quite well (blue predictions overlap the true *F*_ST_ red line in Fig. 6). However, 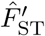 and 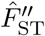 are oracle methods, since they require parameters 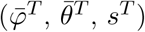 that are not known in practice.

**Figure 6:**
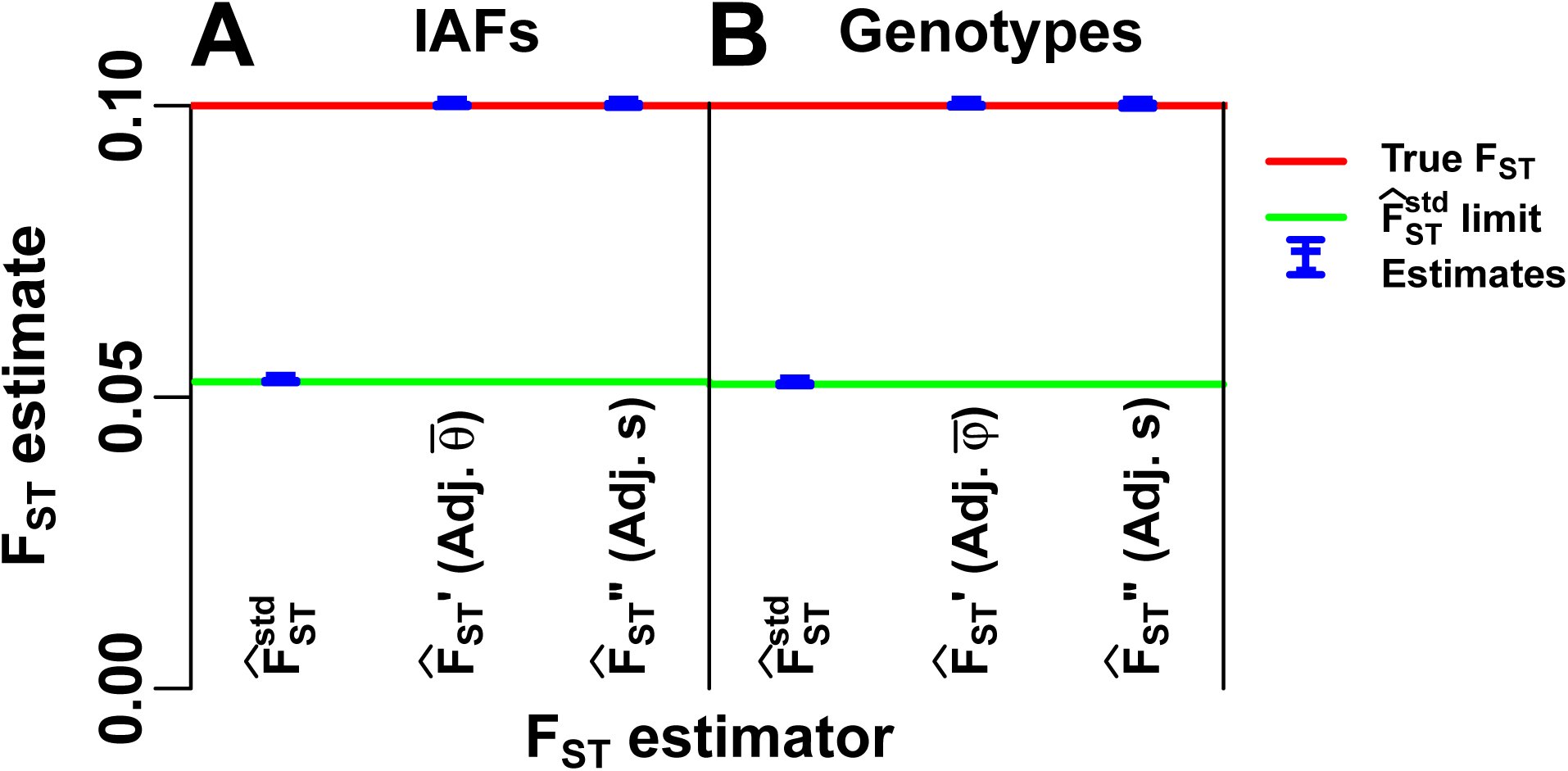
Evaluation of standard and adjusted *F*_ST_ estimators. The convergence values we calculated for the standard kinship plug-in and adjusted *F*_ST_ estimators are validated using our admixture simulation. All adjusted estimators are unbiased but are “oracle” methods, since the mean kinship 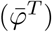, mean coancestry 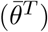, or bias coefficient (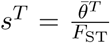 for IAFs, replaced by 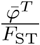 for genotypes) are usually unknown. A) Estimation from individual-specific allele frequencies (IAFs): 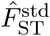 is the standard coancestry plug-in estimator in Eq. (21); 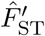“Adj. 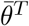” is in Eq. (22); 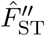“Adj. *s*” is in Eq. (26). B) For genotypes, 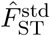 is given in Eq. (20), and the adjusted estimators use 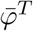 rather than 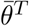. Lines: true *F*_ST_ (red line), limits of biased estimators 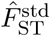 (green lines, which differ slightly per panel). Estimates (blue) include 95% prediction intervals (too narrow to see) from 39 independently-simulated genotype matrices for our admixture model (Supplementary Information, Section S9).

Prediction intervals were computed from estimates over 39 independently-simulated IAF and genotype matrices (Supplementary Information, Section S9). Estimator limits are always contained in these intervals because the number of independent loci (*m* = 300, 000) is sufficiently large. Estimates that use genotypes have wider intervals than estimates from IAFs; however, IAFs are not known in practice, and use of estimated IAFs might increase noise. Genetic linkage, not present in our simulation, will also increase noise in real data.

## 7 Discussion

We studied analytically the most commonly-used estimators of *F*_ST_ and kinship, which can be derived using the method of moments. We determined the bias of these estimators under two models of arbitrary population structure (Fig. 1). We calculated the bias of these *F*_ST_ estimators when the independent subpopulations model assumption is violated. This bias is present even when individual-specific allele frequencies are known without error. We also showed that the standard kinship estimator is biased on structured populations (particularly when the average kinship is comparable to the kinship coefficients of interest), and this bias varies for each pair of individuals. These results led us to a new kinship estimator, which is consistent if the minimum kinship is estimated consistently (Fig. 1). We presented an implementation of this approach, which is practically unbiased in our simulations. Our kinship and *F*_ST_ estimates in human data are consistent with the African Origins model while suggesting that human differentiation is considerably greater than previously estimated [47].

Estimation of *F*_ST_ in the correct scale is crucial for its interpretation as an IBD probability, for obtaining comparable estimates in different datasets and across species, as well as for DNA forensics [5, 9, 55, 61, 63–65]. Our findings that existing genome-wide *F*_ST_ estimates are downwardly biased matters in these settings. However, our findings may not have direct implications for per-locus *F*_ST_ estimate approaches where only the relative ranking matters, such as for the identification of loci under selection [10, 12, 66–71], assuming that the bias of the genome-wide estimator carries over uniformly to all per-locus estimates. Note that our convergence calculations in Section 2 require large numbers of loci so they do not apply to single locus estimates. Moreover, various methods for per-locus *F*_ST_ estimation for multiple alleles suffer from a strong dependence to the maximum allele frequency and heterozygosity [68–70, 72–75] that suggests that a more complicated bias is present in these per-locus *F*_ST_ estimators.

We have shown that the misapplication of existing *F*_ST_ estimators for independent subpopulations may lead to downwardly-biased estimates that can approach zero even when the true generalized *F*_ST_ is large. Weir-Cockerham [19], HudsonK (which generalizes the Hudson pairwise *F*_ST_ estimator [20] to *K* independent populations), and BayeScan [12] *F*_ST_ estimates in our admixture simulation are biased by nearly a factor of two (Fig. 4B), and differ from our new *F*_ST_ estimates in humans by nearly a factor of three [47]. These estimators were derived assuming independent subpopulations, so the observed biases arise from their misapplication to subpopulations that are neither independent not homogeneous. Nevertheless, natural populations—particularly humans— often do not adhere to the independent subpopulations model [47, 76–80] (also see Section 2 in Part I).

The standard kinship coefficient estimator we investigated is often used to control for population structure in GWAS and to estimate genome-wide heritability [18, 24, 27–32]. While this estimator was known to be biased [18, 32], no closed form limit had been calculated until now (concurrently calculated by [53]). We found that kinship estimates are biased downwards on average, but bias also varies for each pair of individuals (Fig. 1, Fig. 5). Thus, the use of these distorted kinship estimates may be problematic in GWAS or for estimating heritability, but the extent of the problem remains to be determined.

We developed a theoretical framework for assessing genome-wide ratio estimators of *F*_ST_ and kinship. We proved that common ratio-of-means estimators converge almost surely to the ratio of expectations for infinite independent loci (Supplementary Information, Section S1.1). Our result justifies approximating the expectation of a ratio-of-means estimator with the ratio of expectations [6, 19, 20]. However, mean-of-ratios estimators may not converge to the ratio of expectations for infinite loci. Mean-of-ratios estimators are potentially asymptotically unbiased for infinite individuals, but it is unclear which estimators have this behavior. We found that the ratio-of-means kinship estimator had much smaller errors from the ratio of expectations than the more common mean-of-ratios estimator, whose convergence value is unknown. Therefore, we recommend ratio-of-means estimators, whose asymptotic behavior is well understood.

We have demonstrated the need for new models and methods to study complex population structures, and have proposed a new approach for kinship and *F*_ST_ estimation that provides nearly unbiased estimates in this setting. Extending our implementation to deliver consistent accuracy in arbitrary population structures will require further innovation, and the results provided here may be useful in leading to more robust estimators in the future.

## Software

An R package called popkin, which implements the kinship and *F*_ST_ estimation methods proposed here, is available on the Comprehensive R Archive Network (CRAN) at https://cran.r-project.org/package=popkin and on GitHub at https://github.com/StoreyLab/popkin.

An R package called bnpsd, which implements the BN-PSD admixture simulation, is available on CRAN at https://cran.r-project.org/package=bnpsd and on GitHub at https://github.com/StoreyLab/bnpsd.

An R package called popkinsuppl, which implements memory-efficient algorithms for the WC and HudsonK *F*_ST_ estimators, and the standard kinship estimator, is available on GitHub at https://github.com/OchoaLab/popkinsuppl.

Public code reproducing these analyses are available at https://github.com/StoreyLab/human-differentiation-manuscript.

## Acknowledgments

This research was supported in part by NIH grant R01 HG006448.

## Supplementary Information

### S1 Accuracy of ratio estimators

#### S1.1 Almost sure convergence of ratio-of-means estimators with independent and uniformly-bounded terms

Here we prove that 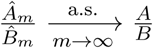, where 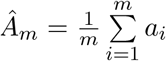 and 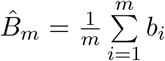 give the ratio-of-means estimator described in the main text. It suffices to prove 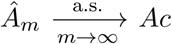 and 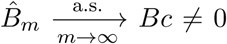 from which the result follows using the continuous mapping theorem [81, 82]. The proof for 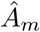 follows, which applies analogously to 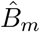. Our *a*_*i*_ are independent but not identically distributed, since they depend on 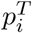 that varies per locus, so the standard law of large numbers does not apply to 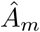. We show almost sure convergence using Kolmogorov’s criterion for the Strong Law of Large Numbers [83], which is satisfied for bounded Var(*a*_*i*_). Since |*a*_*i*_| ≤ *C* < *∞* for all *i* and some *C* (see main text), then 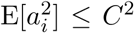, so Var(*a*_*i*_) ≤ *C*^2^. Therefore, 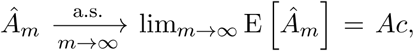, as desired.

#### S1.2 Order of error of expectations

The error of the ratio of expectations from the expectation of the ratio is given by

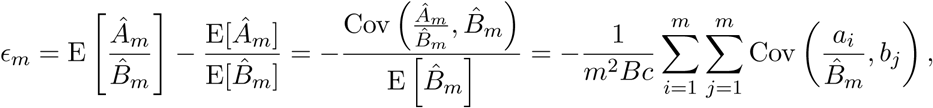

which follows from Cov(*X, Y*) = E[*XY*] − E[*X*] E[*Y*] and expanding the covariance [84]. Previous work on ratio estimators [54, 84] assumes IID *a*_*i*_ and *b*_*i*_, which does not hold for SNP loci. Assuming independent loci (Cov(*a*_*i*_, *b*_*j*_) = 0 for *i* ≠ *j*) and large *m* so 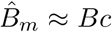 is practically independent of any given *a*_*i*_ and *b*_*j*_, then

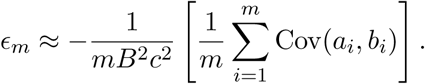

Since *a*_*i*_, *b*_*i*_ are bounded, |Cov(*a*_*i*_, *b*_*i*_)| ≤ *C*^2^ for the same *C* of the previous section, so

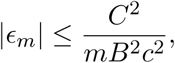

for some large enough *m* and *C*. Hence 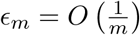 as is for standard ratio estimators [54].

### S2 Previous *F*_ST_ estimators for the independent subpopulations model

Here we summarize the previous WC and Hudson *F*_ST_ estimators for independent subpopulations and introduce the generalized HudsonK estimator for more than two subpopulations. In this section, let *i* index the *m* loci, *j* index the *n* subpopulations, *n*_*j*_ be the number of individuals sampled from subpopulation *j*, and 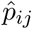 be the sample reference allele frequency at locus *i* in subpopulation *j*.

#### S2.1 The Weir-Cockerham *F*_ST_ estimator

The Weir-Cockerham (WC) *F*_ST_ estimator [19] estimates the coancestry parameter *θ*^*T*^ shared by each of the *n* independent subpopulation in consideration. Let *ĥ*_*ij*_ denote the fraction of heterozygotes in subpopulation *j* for locus *i*. The ratio-of-means WC *F*_ST_ estimator and its limit for independent subpopulations (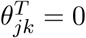 for *j* ≠ *k*) with equal differentiation 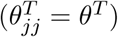 is

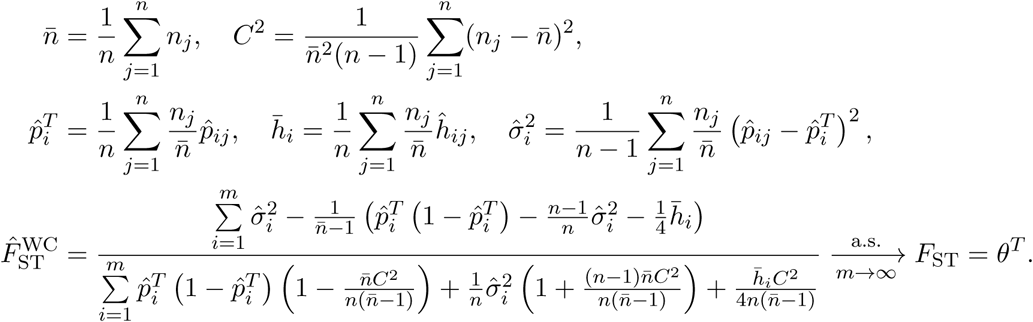

Note that 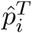 above weighs every individual equally by weighing subpopulation *j* proportional to its sample size *n*_*j*_, so it equals the estimator in Eq. (10) with uniform weights.

Now we simplify this estimator as the sample size of every subpopulation becomes infinite. First set the sample size of every subpopulation *n*_*j*_ equal to their mean 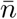, which implies *C*^2^ = 0 and

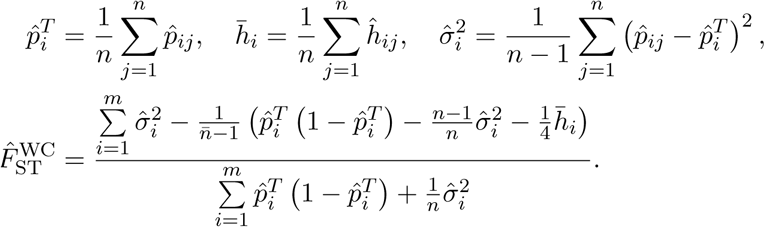

Now we take the limit as the sample size 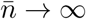, which results in sample allele frequencies converging to the true subpopulation allele frequencies 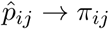 for every subpopulation *j* and locus *i*, and

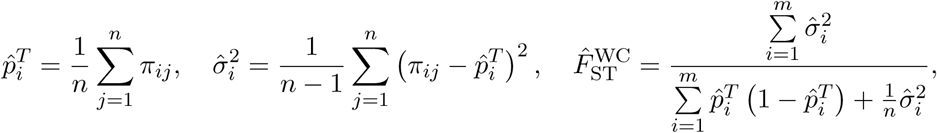

which matches the 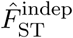 in Eqs. (2) to (4) as desired. Note the number of subpopulations *n* remains finite, and the sample heterozygosity 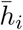 is not needed in the limit.

#### S2.2 The Hudson *F*_ST_ estimator

The Hudson pairwise *F*_ST_ estimator [20] measures the differentiation of two subpopulations (*j, k*). The estimator and its limit for two independent subpopulations 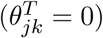 is

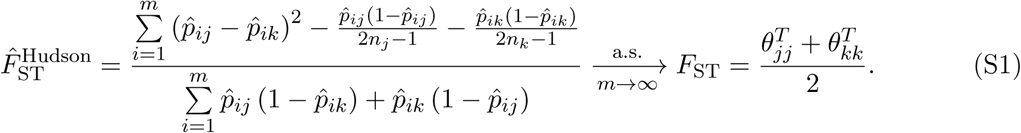

#### S2.3 Generalized HudsonK *F*_ST_ estimator

Here we present the “HudsonK” estimator, which generalizes the Hudson pairwise *F*_ST_ estimator in Eq. (S1) to *n* independent subpopulations. Note that for independent subpopulations, the *F*_ST_ of all the subpopulations equals the mean pairwise *F*_ST_ of every pair of subpopulations:

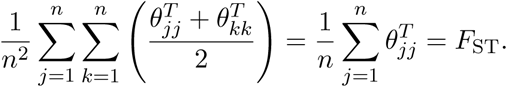

For that reason, averaging numerators and denominators of the pairwise estimator in Eq. (S1) before computing the ratio, we obtain the generalized estimator and a limit under independent subpopulations of

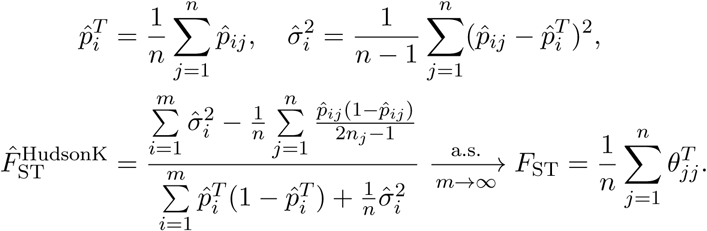

Note that unlike the WC estimator, 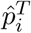 above weighs every subpopulation equally, so every individual is weighed inversely proportional to the sample sizes *n*_*j*_ of their subpopulation *j*.

Like 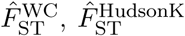 simplifies to 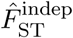 in Eqs. (2) to (4) in the limit of infinite sample sizes *n*_*j*_ → ∞, where 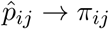 for every (*i, j*).

### S3 Derivation of method-of-moment estimators

#### S3.1 *F*_ST_ estimator for independent subpopulations

Assuming the coancestry model in Eqs. (5) and (6) for independent subpopulations (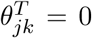 for *j* ≠ *k*), the first and second moments of the IAFs are:

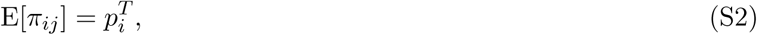

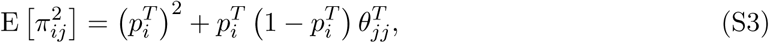

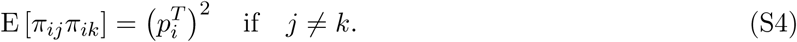

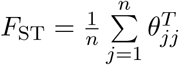 appears by averaging Eq. (S3) over *j*:

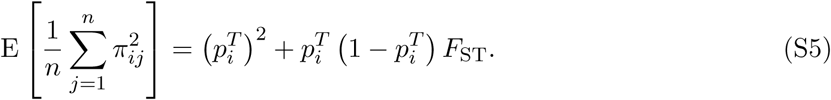

Since Eq. (S2) has the same value for every *j*, and Eq. (S4) as well for every *j* ≠ *k*, we average these to reduce estimation variance. The results are in terms of 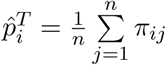:

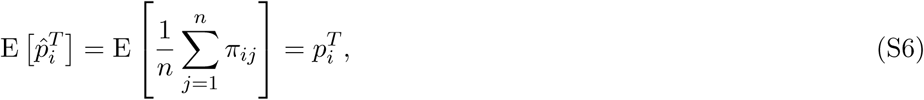

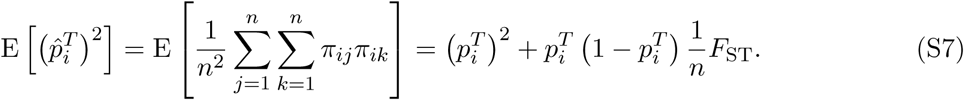

*F*_ST_ also appears in Eq. (S7) because *j* = *k* terms are introduced in the double sum. Subtracting Eq. (S5) and Eq. (S7) in turn from Eq. (S6) results in:

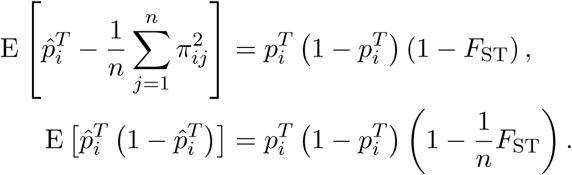

To reduce variance further, we average across loci, giving

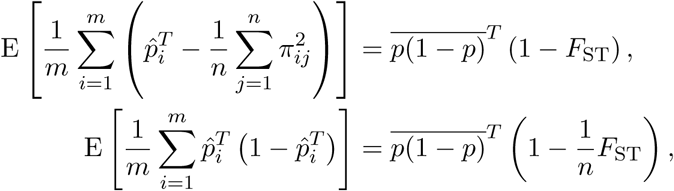

where 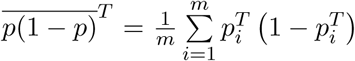. Eliminating 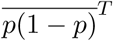 and solving for *F*_ST_ in this system of equations results in the following *F*_ST_ estimator:

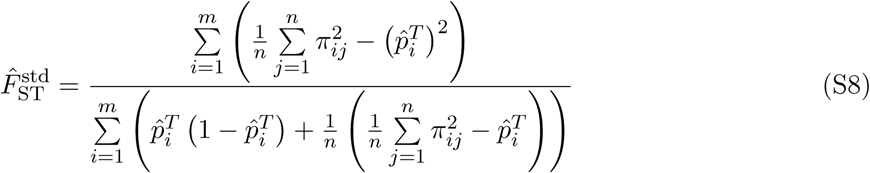

This estimator is simplified noting that 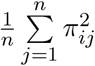 appears in the IAF sample variance,

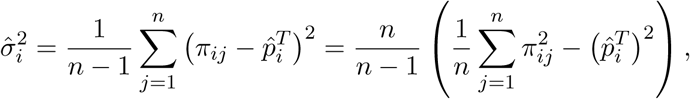

so substituting it into Eq. (S8) recovers Eq. (4) as desired:

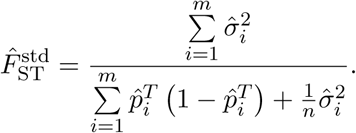

#### S3.2 Standard kinship estimator

Here we assume the kinship model in Eqs. (12) and (13). Since Eq. (12) is the same for all individuals *j*, we average these first moments to reduce variance,

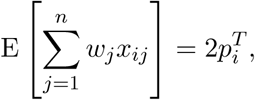

which results in the following estimator of 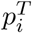:

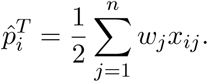

Each 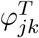 appears once per (*j, k*) pair in Eq. (13), recast here in terms of the sample covariance:

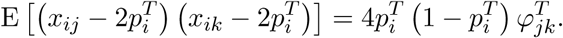

Variance in the kinship estimate is reduced by averaging across loci, yielding:

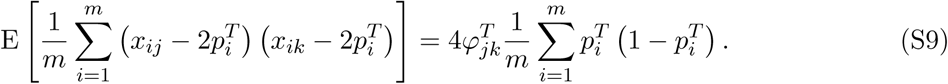

Plugging 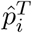 into Eq. (S9) and solving for 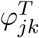 recovers Eq. (11) as desired:

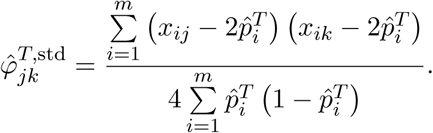

### S4 Proofs that *F*_ST_ and kinship estimator limits are constants with respect to the ancestral population *T*

In our work we calculate the limits of several estimators, which are given in terms of an arbitrary ancestral population *T* (not necessarily the MRCA, unless otherwise noted). The apparent paradox that the limit of an estimator would vary depending on the choice of *T* is resolved since these limits are in fact constant with respect to *T*. All proofs depend on the following IBD identities for change of ancestral population (see Section 3.4 of Part I for details):

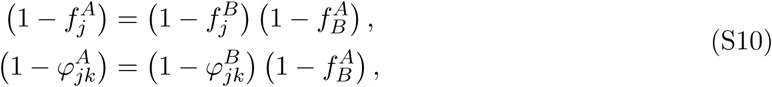

where *A, B* are two possible ancestral populations for the individuals *j, k*, and *A* is ancestral to *B*.

#### S4.1 Proof that the limit of 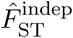 does not depend on *T*

Here we study the limit of 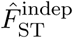 in Eq. (7). Let *S* be a reference population ancestral to the individuals in question and *T* be another population ancestral to *S*. Denote the key parameters relative to *S* by 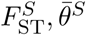 and relative to *T* by 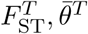. The equations that relate both quantities satisfy our IBD shift identity (which follows by averaging Eq. (S10) over individuals for *F*_ST_ or pairs of individuals for 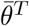):

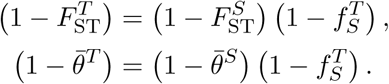

Solving for the values relative to *S* gives

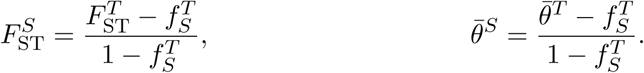

The desired equality of the limit for both *S* and *T* follows:

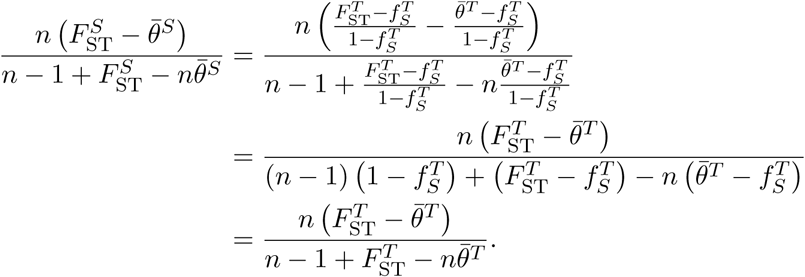

#### S4.2 Proof that the limit of 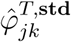 does not depend on *T*

Here we study the limit of the standard kinship estimator 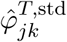 in Eq. (14). Let *S* be a reference population ancestral to the individuals in question and *T* be another population ancestral to *S*. The equations that relate the terms relative to *S* and those relative to *T* follow from Eq. (S10) just as in the previous subsection:

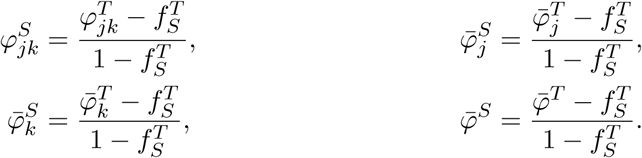

The desired result follows:

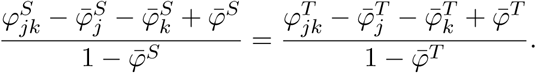

### S5 Mean coancestry bounds

Here we prove that, for any weights such that 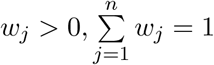,

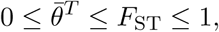

and for uniform weights 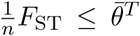. Furthermore, 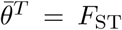 iff 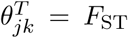 for all (*j, k*), and 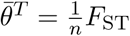 for the independent subpopulations model.

The Cauchy-Schwarz inequality for covariances implies 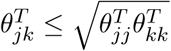. Therefore,

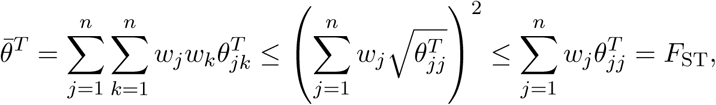

where the second inequality follows from Jensen’s inequality, since *x*^2^ is a convex function. Since 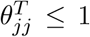, then *F*_ST_ ≤ 1 as well. Equality in the second bound requires 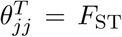 for all *j*, and equality in the first bound requires 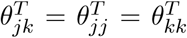, so that 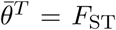 requires 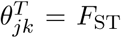 for all (*j, k*). Since all *w*_*j*_, 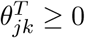, then

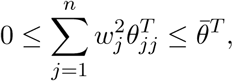

where the second inequality follows from dropping *j* ≠ *k* terms from the double sum of 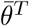. The case 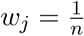 gives 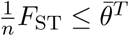, with equality for the independent subpopulations model by construction.

### S6 Moments of estimator building blocks

Here we calculate first and some second moments for “building block” quantities that recur in our estimators, particularly terms involving *x*_*ij*_ and 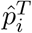, and which enable us to calculate the limits of our estimators. Below are examples for genotypes, which follow from Eqs. (12) and (13); calculations for IAFs follow analogously from Eqs. (5) and (6) (not shown).

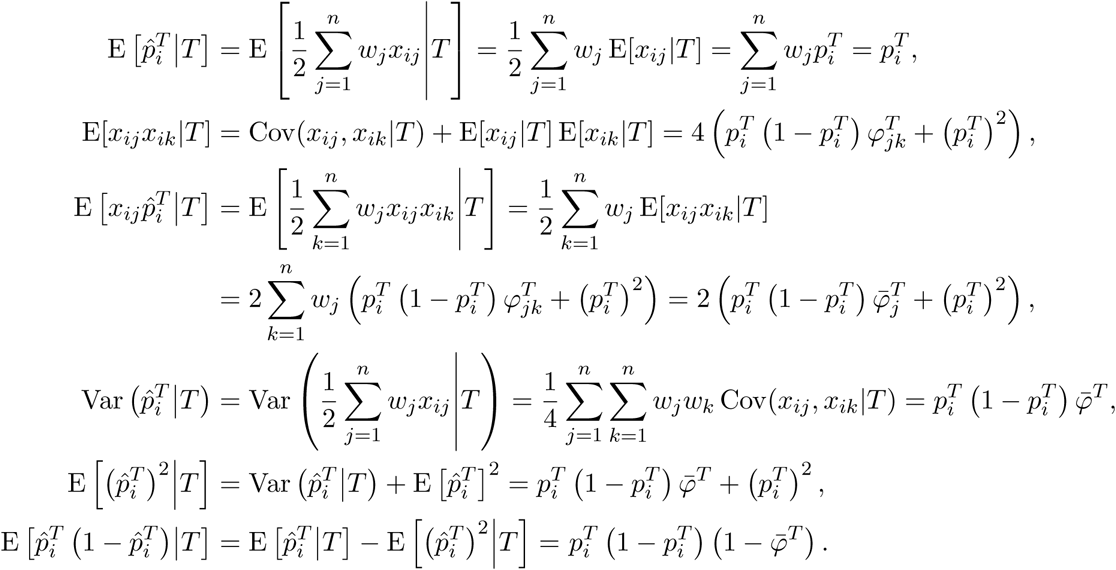

### S7 Derivation of new kinship estimator

To begin the method-of-moments derivation, we compute the raw first and second moments from the kinship model of Eqs. (12) and (13).

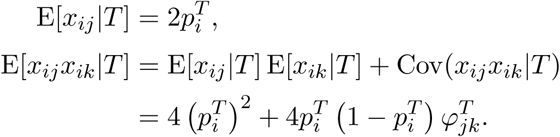

For obtain a symmetric estimator, we also compute the raw moments of 2 − *x*_*ij*_ (which counts the alternative allele):

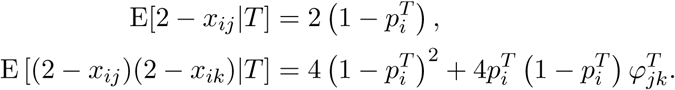

If we solved for 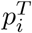 using the first moment equations, we would recover the standard kinship estimator of Eqs. (10) and (11), so we shall avoid this strategy.

To proceed, we average the two second moment equations above. Note that

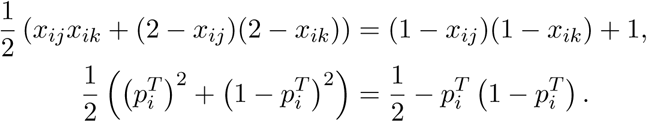

Therefore, the symmetric estimator (which gives the same calculation if the reference allele is switched) is

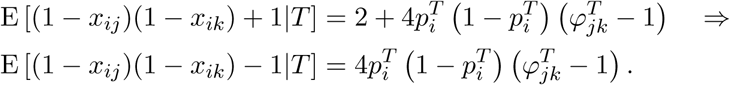

A genome-wide estimate is obtained by averaging the previous statistics across loci, resulting in

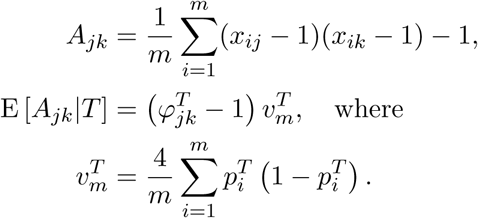

The new kinship estimator follows from obtaining a consistent estimator of the limit of 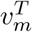 as *m* goes to infinity, and applying it to solve for 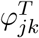 in the above equation for the expectation of *A*_*jk*_, as detailed in Section 5.

### S8 Admixture and independent subpopulations model simulations

#### S8.1 Construction of subpopulation allele frequencies

We simulate *K* = 10 subpopulations *S*_*u*_ and *m* = 300, 000 independent loci. Every locus *i* draws 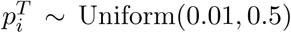. We set 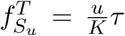 where *τ* ≤ 1 tunes *F*_ST_. For the independent sub-populations model, 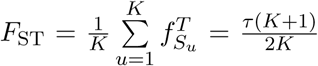, so 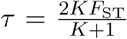 gives the desired *F*_ST_ (*τ* ≈ 0.18 for *F*_ST_ = 0.1). For the admixture model, *τ* is found numerically (*τ* ≈ 0.90 for *F*_ST_ = 0.1; see last subsection). Lastly, 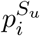 values are drawn from the Balding-Nichols distribution,

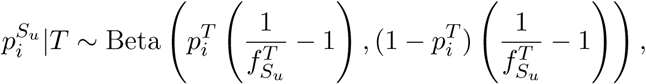

which results in subpopulation allele frequencies that obey the coancestry model of Eqs. (5) and (6), with 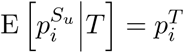 and 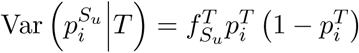 [5], as desired.

#### S8.2 Random subpopulation sizes

We randomly generate sample sizes **r** = (*r*_*u*_) for *K* subpopulations and 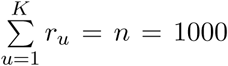 individuals, as follows. First, draw **x** ∼ Dirichlet (1,…, 1) of length *K* and **r** = round(*n***x**). While 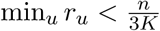, draw a new **r**, to prevent small subpopulations (they do not occur in real data). Due to rounding, 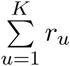 may not equal *n* as desired. Thus, while 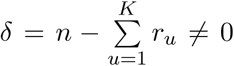, a random *u* is updated to *r*_*u*_ ← *r*_*u*_ + sgn(*δ*), which brings *δ* closer to zero at every iteration. Weights for individuals *j* in *S*_*u*_ are 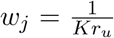 so the generalized *F*_ST_ matches 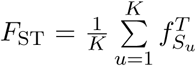 from the independent subpopulations model (Section 3.3.2 of Part I), which HudsonK estimates.

#### S8.3 Admixture proportions from 1D geography

We construct *q*_*ju*_ from random-walk migrations along a one-dimensional geography. Let *x*_*u*_ be the coordinate of intermediate subpopulation *u* and *y*_*j*_ the coordinate of a modern individual *j*. We assume *q*_*ju*_ is proportional to *f* (|*x*_*u*_ − *y*_*j*_|), or

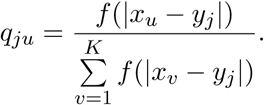

where *f* is the Normal density function with *µ* = 0 and tunable *σ*. The Normal density models random walks, where *σ* sets the spread of the populations (Fig. 5). Our simulation uses *x*_*u*_ = *u* and 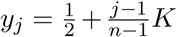, so the intermediate subpopulations span [1, *K*] and individuals span 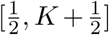. For the *F*_ST_ estimators that require subpopulations, individual *j* is assigned to the nearest subpopulation *S*_*u*_ (the *u* that minimizes |*x*_*u*_ − *y*_*j*_|; Fig. 3D); these subpopulations have equal sample size, so 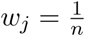 is appropriate.

#### S8.4 Choosing *σ* and *τ*

Here we find values for *σ* (controls *q*_*jk*_) and *τ* (scales 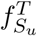) that give 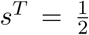 and *F*_ST_ = 0.1 in the admixture model. We previously found that 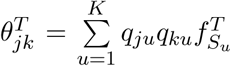 and 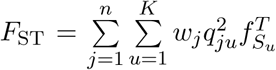 for the BN-PSD model (Section 6.1 of Part I). In our simulation, 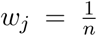 and 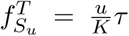, so 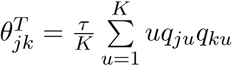 and 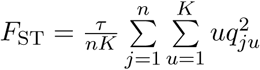. Therefore,

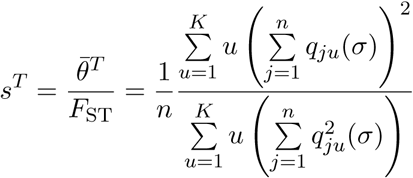

depends only on *σ*. A numerical root finder finds that *σ* ≈ 1.78 gives 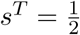. For fixed *q*_*ju*_,

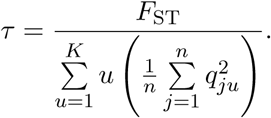

*F*_ST_ = 0.1 is achieved with *τ* ≈ 0.901.

### S9 Prediction intervals of *F*_ST_ estimators

Prediction intervals with *α* = 95% correspond to the range of *n* = 39 independent *F*_ST_ estimates. In the general case, *n* independent statistics are given in order *X*_(1)_ <…< *X*_(*n*)_. Then *I* = [*X*_(*j*)_, *X*_(*n*+1*−j*)_] is a prediction interval with confidence 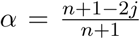 [85]. In our case, *j* = 1 and *n* = 39 gives *α* = 0.95, as desired. Each estimate was constructed from simulated data with the same dimensions and structure as before (fixed 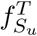 and *q*_*ju*_; fixed sample sizes for the independent subpopulations model), but with 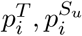, *π*_*ij*_, *x*_*ij*_ drawn separately for each estimate.

## Notes

https://github.com/StoreyLab/human-differentiation-manuscript

